# Differential locus coeruleus–hippocampus interactions during offline states

**DOI:** 10.1101/2025.09.18.677005

**Authors:** Mingyu Yang, Oxana Eschenko

## Abstract

Patterns of locus coeruleus (LC) activity and norepinephrine (NE) release during non-rapid-eye-movement (NREM) sleep suggest a critical role for the LC–NE system in offline modulation of forebrain circuits. NE transmission promotes synaptic plasticity and is required for memory consolidation, but the field has only begun to uncover how LC activity contributes to coordinated forebrain network dynamics. Hippocampal ripples, a hallmark of memory replay, are temporally coupled with thalamocortical oscillations; however, the circuit mechanisms underlying systems-level consolidation across larger brain networks remain incompletely understood. Here, using multi-site electrophysiology, we examined LC firing in relation to hippocampal ripples in freely behaving rats. LC activity and ripple occurrence were state-dependent and inversely related: heightened arousal was associated with increased LC firing and reduced ripple rates. At finer timescales, LC spiking decreased ∼1–2 seconds before ripple onset, with the strongest modulation during awake ripples but minimal change during ripple–spindle coupling. These findings reveal state-dependent dynamics of LC-hippocampal interactions, positioning the LC as a key component of a cortical–subcortical network supporting systems-level memory consolidation.

## Introduction

Norepinephrine (NE) modulation of the forebrain circuits supporting cognition has long been considered to play a role during vigilant states (***Sara, 2009***; ***Sara and Bouret, 2012***), whereas its impact during low arousal (or ‘offline’) states, including sleep, has received less attention. The pioneering discovery of greatly reduced firing of the locus coeruleus (LC) neurons during sleep (***Aston-Jones and Bloom, 1981***), for decades, directed research focus toward the role of LC, as a part of the ascending arousal system, for ‘online’ information processing. At the same time, pharmacological studies in both animals (***Sara et al., 1999***; ***Roullet and Sara, 1998***; ***Clayton and Williams, 2000***; ***Galeotti et al., 2004***; ***Gazarini et al., 2013***) and humans (***Groch et al., 2011***; ***Gais et al., 2011***; ***Kuffel et al., 2014***; ***Cahill et al., 1994***) have demonstrated the importance of post-learning NE transmission for memory consolidation. The results of pharmacology studies were consistent with a well-established facilitatory role of NE for synaptic plasticity, which also takes place offline (***Straube and Frey, 2003***; ***Gelinas and Nguyen, 2005***; ***Harley, 2007***; ***Hansen and Manahan-Vaughan, 2015***; ***Hagena et al., 2016***; ***Palacios-Filardo and Mellor, 2019***).

During offline states such as awake immobility and non-rapid eye movement (NREM) sleep, hippocampal activity is marked by transient episodes of synchronized firing, detected in CA1 local field potentials (LFPs) as high-frequency ∼150 Hz oscillations known as ripples **Buzsaki** (**1996**). Hippocampal ripples, a hallmark of memory replay, are temporally coupled with thalamocortical sleep spindles (10–16 Hz) and cortical slow (∼1Hz) oscillations, which together represent key components of the mechanism underlying systems-level memory consolidation (***Klinzing et al., 2019***). In recent years, it became evident that ripples reflect not only a coordinated local activity within the hippocampus but also indicate the emergence of large-scale functional networks (***Logothetis et al., 2012***; ***Nitzan et al., 2022***). The ripple-associated cross-regional communication may occur through a coordinated up/down-regulation of neural activity across cortical and subcortical structures, including neuromodulatory centers (***Skelin et al., 2018***; ***Maingret et al., 2016***; ***Brodt et al., 2023***). Several studies have demonstrated temporally coordinated cross-regional neuronal activity around ripples, involving the amygdala (***Girardeau et al., 2017***), the ventral tegmental area (***Gomperts et al., 2015***), the median raphe (***Wang et al., 2015***), and the thalamus (***Logothetis, 2015***; ***Yang et al., 2019***; ***Varela et al., 2001***).

While some indirect evidence suggests a link (***Logothetis et al., 2012***), the relationship between LC neuron spiking and hippocampal ripples has not been directly characterized. For example, a study in the murine slice showed that increasing NE concentration increased ripple incidence and their amplitudes (***Ul Haq et al., 2012***). Consistently, in behaving rats, we showed that pharmacological suppression of NE transmission decreased the occurrence of ripples and impaired spatial memory consolidation (***Duran et al., 2023***). Finally, experimentally induced LC activation affected the temporal pattern of ripple occurrence (***Novitskaya et al., 2016***; ***Swift et al., 2018***). Earlier studies have shown close temporal relations between LC and sleep spindles (***Aston-Jones and Bloom, 1981***; ***Swift et al., 2018***; ***Osorio-Forero et al., 2021***; ***Kjaerby et al., 2022***). In our previous work, we have shown that firing of a subpopulation of LC neurons was temporally coordinated with slow oscillations (***Eschenko et al., 2012***; ***Totah et al., 2018***), which in turn orchestrate spindle and ripple activity (***Molle et al., 2006***).

In the present study, we characterized the ripple-associated LC firing patterns in freely behaving rats. We report that LC activity was anticorrelated with the ripple rate at both slow (multi-second) and fast (sub-second) timescales. Our results also provide evidence for a state-dependent engagement of the LC in offline memory processing and hippocampal-cortical communication.

## Results

We analyzed a total of 20 recording sessions (n = 7 rats, 2 to 8 sessions per rat with an average duration of 6458.48 ± 190.43 sec), including two datasets (n = 1 rat) that contributed to our previous publication (***Eschenko et al., 2012***). All datasets contained simultaneously recorded extracellular spikes of putative LC-NE neurons, local field potentials (LFPs) from the CA1 subfield of the dorsal hippocampus (HPC), and frontal EEG (Figure 1). The recordings were performed in freely behaving adult male rats placed in a small chamber (30 x 30 x 40 cm) in dim light during the dark phase of the animal’s circadian rhythm. Electrode positioning was fine-tuned using movable microdrives to ensure optimal detection of hippocampal ripples and LC spikes. The recordings from putative LC-NE neurons were verified using multiple electrophysiological criteria (see Methods for details), and the recording sites were additionally confirmed by histological examination (Figure 1B). The average firing rate of LC single units was 1.70 ± 0.21 Hz during wakefulness, 0.51 ± 0.07 Hz during NREM sleep, and 0.014 ± 0.01 Hz during REM sleep, differing significantly across arousal states (one-way ANOVA: F(2,38) = 39.8, *p* < 0.0001; Figure 1-figure supplement 1). This firing pattern is characteristic of LC-NE neurons and is consistent with existing literature (***Aston-Jones and Bloom, 1981***; ***Eschenko et al., 2012***; ***Eschenko and Sara, 2008***; ***Hayat et al., 2020***; ***Takahashi et al., 2010***).

**Figure 1.**
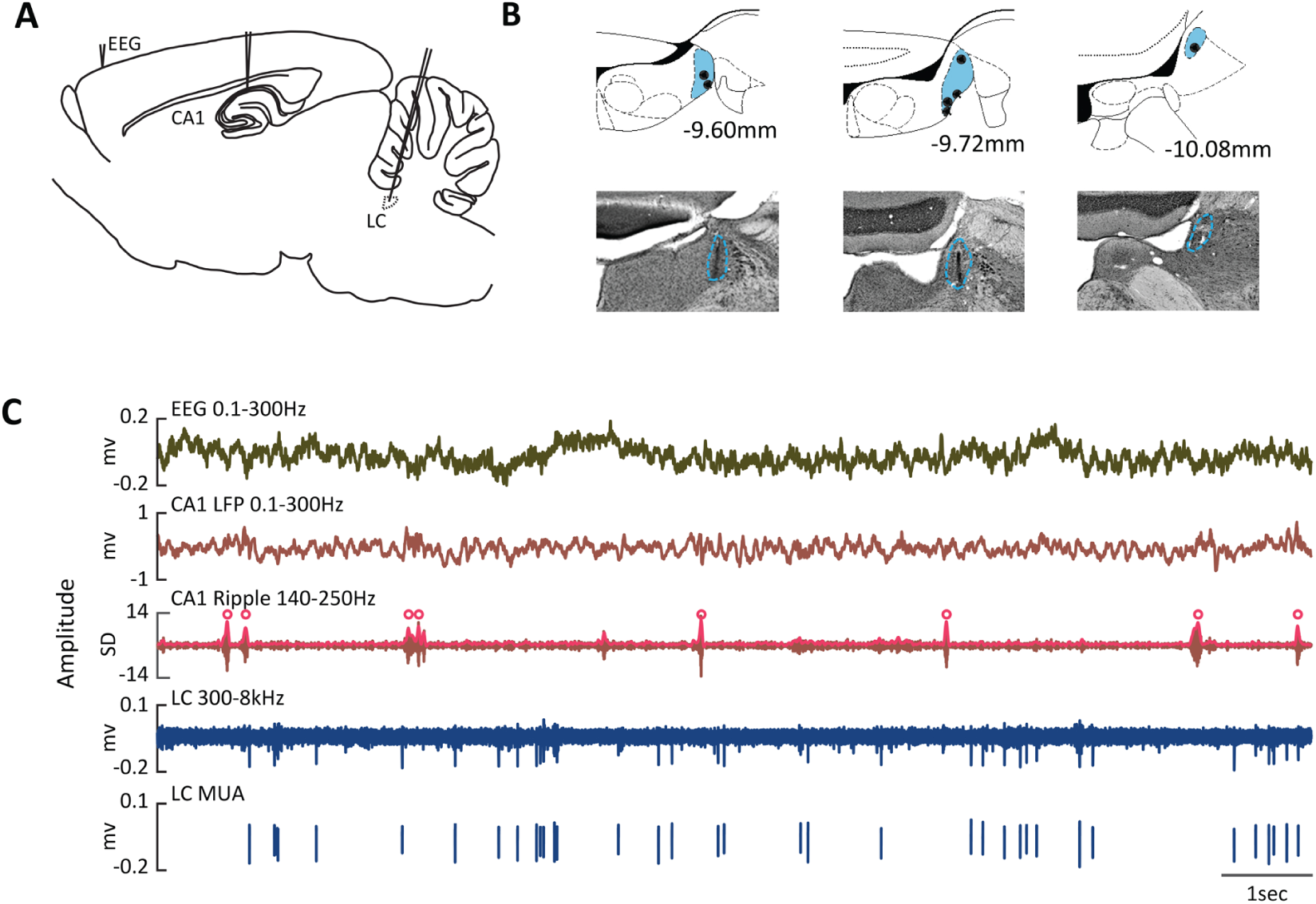
Multi-site extracellular recordings in freely behaving rats. **(A)** Schematic of chronically implanted electrodes for recording the frontal EEG, local field potentials (LFPs) in the dorsal hippocampus (CA1), and neuron spiking in the locus coeruleus (LC). **(B)** Histological verification and reconstruction of electrode placements (black dots) within the LC (blue area). Numbers indicate anterior–posterior coordinates relative to bregma (***Paxinos and Watson, 2005***). **(C)** Representative traces of simultaneously recorded EEG, hippocampal LFPs, and LC neuron spikes during quiet awake. Red dots indicate hippocampal ripples.

### LC-NE neuron spiking is suppressed around hippocampal ripples

We first characterized the temporal relationships between the LC neuron spiking and ripple occurrence. Ripples were detected from a power envelope of a band-passed (140 - 250 Hz), smoothed (at 25 Hz), and z-score normalized CA1 LFPs (Figure 1C). The average ripple rate was 18.2 ± 1.5 ripples per minute, with a ripple amplitude of 10.3 ± 0.3 standard deviations (SD). There was a robust CA1 LFP power increase in the ripple band (140 - 250 Hz) around peaks of detected ripples (Figure 2A) and a transient power increase in the EEG delta (1 - 4 Hz) and spindle (12 - 16 Hz) bands (Figure 2B) immediately after the ripple onset, most likely reflecting the well-known temporal coupling of ripples with slow waves and sleep spindles (***Molle et al., 2006***).

**Figure 2.**
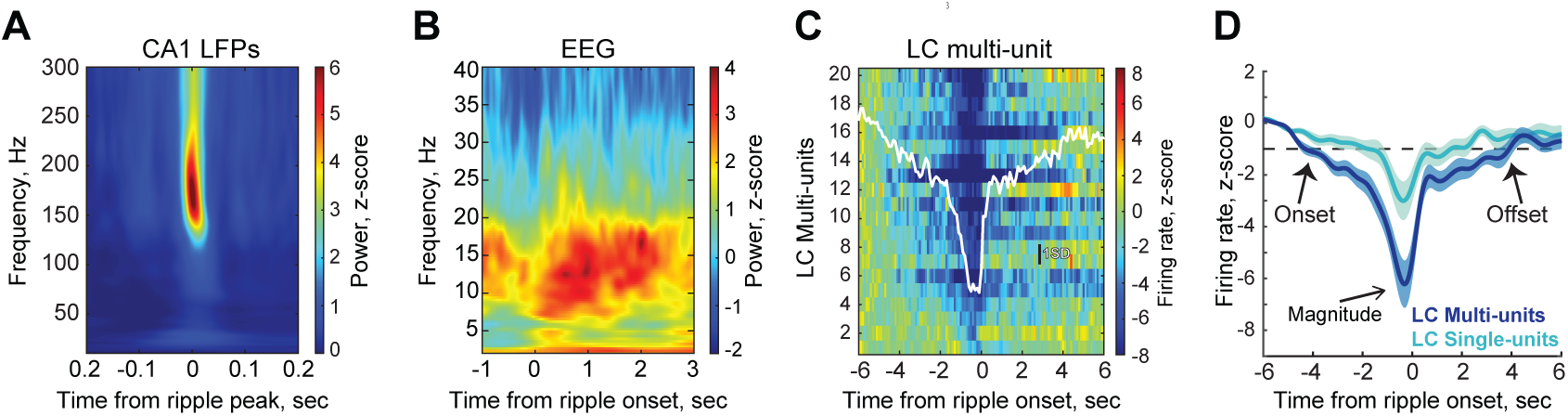
LC-NE neuron spiking is suppressed around hippocampal ripples. (A, **B)** Representative peri-ripple spectrograms of CA1 LFPs **(A)** and frontal EEG **(B)**. **(C)** Normalized peri-ripple LC multi-unit activity. Each row represents an individual dataset, with the overlaid trace showing the average across sessions. **(D)** Comparison of peri-ripple LC multi- and single-unit firing. Normalized firing rates were averaged and smoothed with a 1 Hz low-pass filter. The dashed line marks the 1 SD threshold used to define the onset and offset of ripple-associated LC modulation. The modulation magnitude was extracted as a through on the peri-event histogram.

The LC multiunit activity (LC-MUA) was extracted by thresholding a high-pass filtered (300 Hz –8 kHz) extracellular signal recorded from the LC (Figure 1C). To directly compare the ripple-related LC activity dynamics, for each recording session, we computed the LC-MUA firing rate in a [-6, 6] sec window centered on the ripple onset, z-scored, and averaged across all detected ripples. This analysis revealed a prolonged and consistent suppression of LC activity both before and after ripple onset, with peak suppression < -2 SDs in all sessions (Figure 2C). In six out of twenty recording sessions, we could reliably isolate spikes from a total of 15 single units (n = 4 rats). The peri-ripple (± 6 s) firing suppression of LC single-unit activity (LC-SUA) exceeding the significance threshold of 2 SDs was observed in 13 of 15 cases (Figure 2D).

To quantify the peri-ripple LC activity change, we extracted the modulation onset time, duration, and magnitude, as illustrated in Figure 2D. The modulation magnitude, as measured from the peri-event histogram, was greater for LC-MUA (-6.83 ± 0.84 SDs) compared to LC-SUA (-4.01 ± 0.65 SDs). Although the modulation magnitude differed (Kolmogorov–Smirnov test, p = 0.019), the temporal profiles of both LC-MUA and LC-SUA around ripples were remarkably similar (Figure 2D). No significant differences were found in the onset of spiking suppression (MUA: -4.08 ± 0.71 sec vs. SUA: -2.90 ± 0.50 sec, p = 0.138), duration (MUA: 8.28 ± 1.37 sec vs. SUA: 5.16 ± 0.92 sec, p = 0.069), or peak time (MUA: -0.29 ± 0.04 sec vs. SUA: -0.35 ± 0.04 sec, p = 0.561). Together, these findings indicated that the majority of LC-NE neurons showed a stereotypic activity pattern around hippocampal ripples. Therefore, we used the LC-MUA datasets for subsequent analyses.

### LC firing and ripple occurrence are state-dependent and inversely related

The LC activity around rippled was modulated at multiple temporal scales. In the previous section, we described a relatively sharp drop in the LC firing rate ∼ 2 s before the ripple onset. To capture LC dynamics at longer temporal scales, we tested a range of time windows and found that a 12-s peri-event interval adequately represents LC activity dynamics. When computing peri-ripple LC firing over a [–12, 12] sec time window, we observed a decrease in LC activity beginning as early as ∼10 s before the ripple onset (Figure 2-figure supplement 1). We hypothesized that slower LC dynamics might be related to fluctuations of the global brain state. Indeed, transient spectral changes in the EEG coincided with the occurrence of hippocampal ripples (Figure 2B).

We thus examined the temporal relationships between the ripple and LC activity and ongoing cortical state. To this end, we computed the ripple and LC-MUA rates within 4-s windows and quantified the corresponding cortical state using a synchronization index (SI), calculated as a power ratio (1–4 Hz/30–90 Hz) of the frontal EEG. Figure 3A illustrates that a higher SI corresponded to a lower arousal state; this brain state was associated with more synchronized cortical population activity, a higher ripple rate, and reduced LC neuron firing. Across all sessions, SI values were negatively correlated with LC-MUA (Pearson correlation, r = –0.71 to –0.25, p < 0.01, the highest p = 0.0095; Figure 3B, upper panel) and positively correlated with the ripple rate (r = 0.10 to 0.50, p < 0.01, the highest p = 0.008). As shown in Figure 3B, LC activity was negatively correlated with both the SI and ripple rate (r = –0.38 to –0.11, p < 0.01, with the highest p = 0.0074, n = 20; Figure 3B, lower panel). Together, these results demonstrate that LC activity, cortical state, and ripple occurrence are strongly interrelated, suggesting that the slow (multi-second) dynamics of peri-ripple LC modulation may, in part, reflect underlying brain-state fluctuations.

**Figure 3.**
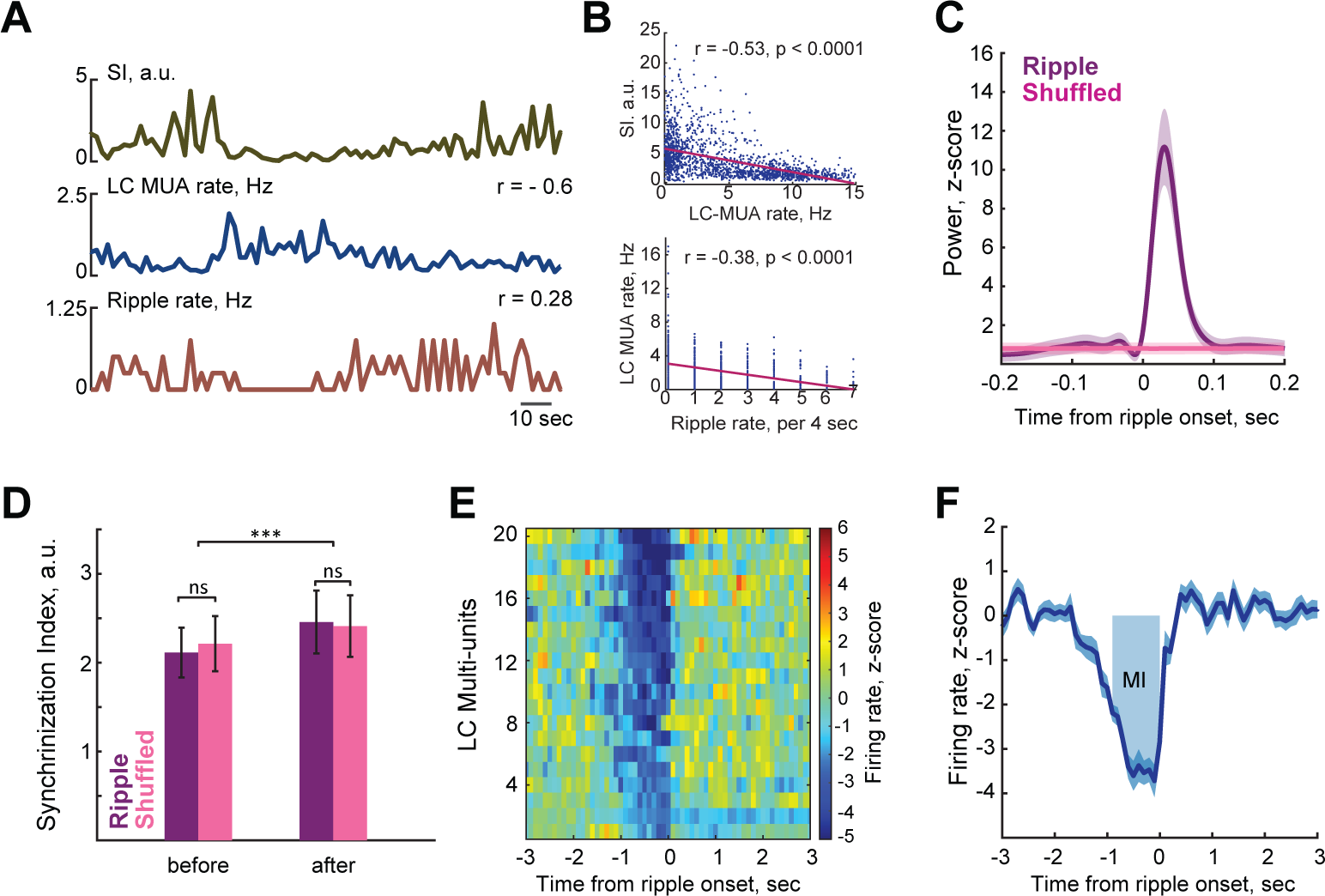
The relationship between LC activity and hippocampal ripples at multiple temporal scales. **(A)** Representative traces showing the relationship among cortical state, LC neuron spiking, and ripple rate. Cortical state was quantified by the Synchronization Index (SI), calculated as the ratio of the EEG delta (1–4 Hz) to gamma (30–90 Hz) power. Higher SI values, reflecting a more synchronized cortical state, were associated with lower LC activity and higher ripple rates. **(B)** An example of correlation between LC-MUA and SI (upper panel) and between LC-MUA and ripple rate (lower panel) obtained from the same recording session. **(C)** Mean CA1 LFPs ripple-band (140–250 Hz) power aligned to the ripple onset (purple) or shuffled events (pink). **(D)** Average Synchronization Index (SI) around ripples and shuffled events. The cortical state preceding shuffled events and ripples was comparable, as confirmed by the absence of significant differences in SI (Wilcoxon signed-rank test; shuffled: Z = -0.20, p = 0.84; ripples: Z = 0.14, p = 0.88). Cortical synchrony increased following both events (shuffled: Z = -3.50, p = 0.00044; ripples: Z = -3.66, p = 0.00026). Similar cortical state dynamics surrounding shuffled events and ripples indicate that the surrogate events adequately capture the cortical state associated with ripple occurrence. **(E)** Normalized peri-ripple LC-MUA for individual sessions. **(F)** Session-averaged peri-ripple LC-MUA. The shaded area denotes the [-1, 0 sec] time window used for the Modulation Index (MI) calculation.

To account for the potential influence of global brain state on peri-ripple LC activity, we generated surrogate events for each session by jittering the timestamps of detected ripples (see Methods). We first verified that hippocampal CA1 LFPs (140–250 Hz) triggered on these surrogate events lacked the ripple-specific frequency component (Figure 3C) and that the shuffled events adequately captured the cortical state dynamics associated with ripples (Figure 3D). Thus, LC activity aligned to surrogate events captured its state-dependent dynamics. To isolate peri-ripple LC modulation from state-dependent fluctuations of the LC activity, we subtracted the LC-MUA around shuffled events from the peri-ripple LC-MUA for each session. The resulting traces provided a state-corrected estimate of ripple-associated LC activity, largely free from confounding effects of cortical state transitions. This analysis revealed a consistent suppression of LC activity at a finer temporal scale (1–2 s), although the temporal profile and magnitude of modulation varied across sessions (Figure 3E).

To quantify the degree of peri-ripple LC activity modulation, we extracted the onset time and the duration as illustrated in Figure 2D. In addition, we calculated the modulation index (MI) as the area under the curve of the session-averaged peri-ripple LC rate within 1 second preceding the ripple onset (Figure 3F, see Methods for details). Because high-quality recordings from the LC in behaving rats are technically challenging and relatively rare, we included all valid sessions in this study. The average MI, calculated per animal and per session, fell within a consistent range despite variability in the number of recording sessions (2–8 sessions per rat) across (Figure 3-figure supplement 1). On average, LC activity significantly decreased 1.41 ± 0.06 sec (range: 1.02 – 1.88 sec) before the ripple onset and returned to the baseline after 1.70 ± 0.08 sec (range: 1.10 – 2.58 sec). Notably, maximal LC activity suppression occurred approximately 200 ms before the ripple onset (0.24 ± 0.04 sec; range: 0.03 – 0.73 sec). For all 20 datasets, the MI exceeded a 95% confidence interval (CI) of the MIs calculated around the shuffled time series, confirming a substantial decrease in LC activity around ripples.

### Differential LC modulation across ripple subsets

The overall transient suppression of LC activity around hippocampal ripples, reported above, does not preclude differential LC modulation depending on ripple subtype. We therefore examined potential heterogeneity in LC modulation within a ± 6-sec time window around ripples. To this end, we randomly selected 20% of ripples from each dataset, calculated the subset modulation index (subMI) for this subset, and repeated the procedure to generate 5000 ripple subsets (537.6 ± 53.0 ripples per subset). SubMI values showed substantial variability across subsets (range: –45.70 to 1.38 a.u.; Figure 4A). On average, more than half of the subsets (65.99 ± 7.08%) exhibited significant peri-ripple LC modulation (subMI >95% CI), whereas the change of LC activity in the remaining subsets did not exceed 95% CI. Importantly, ripple intrinsic properties did not account for this variability: no significant differences were found in peak amplitude (Wilcoxon signed-rank test, Z = 0.63, p = 0.53), duration (Z = –0.52, p = 0.60), or ripple oscillation frequency (Z = 1.47, p = 0.14) between ripples in the 10th and 90th percentiles of the subMI distribution.

**Figure 4.**
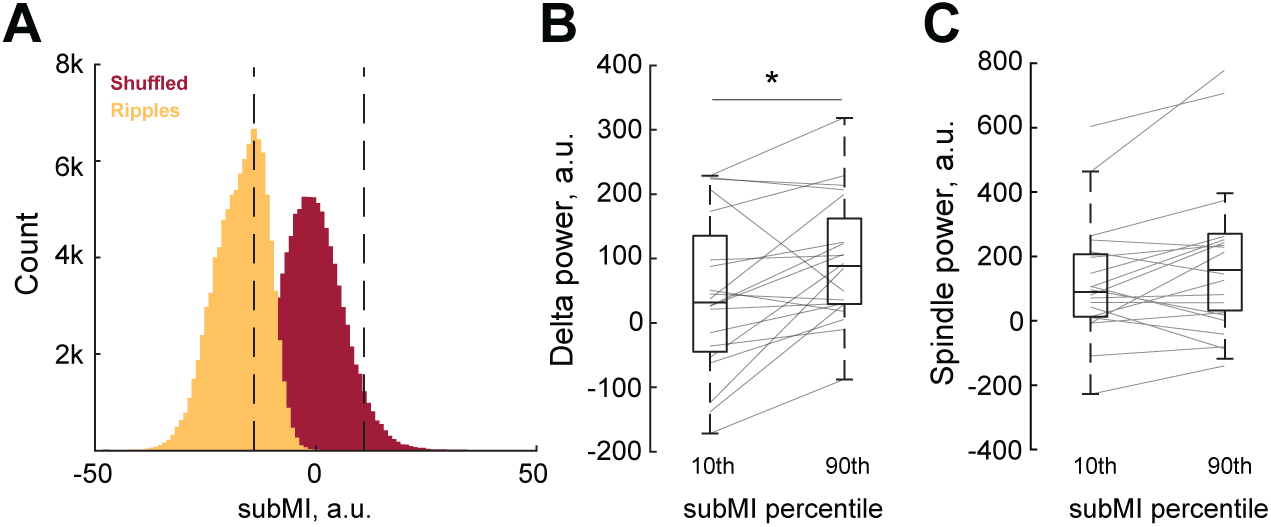
Differential LC modulation across ripple subsets. **(A)** Distribution of modulation index (MI) values for different subsets of ripples (subMI). SubMIs were computed from the peri-event histograms of LC-MUA aligned to the ripple peak (yellow) or ‘shuffled’ time series (red). Shuffled time series were generated by permutation of the inter-ripple intervals; shuffled subMIs were calculated, and the permutation procedure was repeated 5000 times. Vertical dashed lines indicate the 95% confidence interval (CI) boundaries for the shuffled time series; the CI served as the significance threshold. To quantify the LC-MUA modulation around subsets of ripples, subsampled MI (subMI) was computed for a subset of ripples (20% of all detected ripples in each session), and the same procedure was repeated 5000 times (see Methods). **(B, C)** The EEG delta **(B)** and spindle **(C)** power preceding 1 sec the ripple onset plotted for ripple subsets associated with different degrees of LC modulation. Box-whisker plots show the median, the 1st and 3rd quartiles, and min/max for the 10th and 90th percentiles of the subMI distribution, associated with maximal and minimal LC modulation, respectively. Gray lines show data from individual rats. * p < .05 (Wilcoxon signed-rank test). Note a higher EEG delta power preceding ripples, associated with weak or absent LC modulation.

Instead, differences in the ongoing cortical state appeared to underlie the variability in LC modulation. The EEG preceding ripples, which were not associated with LC modulation, had a significantly higher delta (1-4 Hz) power (Wilcoxon signed-rank test, Z = –2.576, p = 0.01; Figures 4B) and enhanced spindle (10-16 Hz) power (Z = –1.829, p = 0.0674; Figures 4C), both indicating a more synchronized cortical state.

### Peri-ripple LC modulation depends on the cortical-hippocampal interaction

Building on our findings, we predicted that the LC-ripple interactions might vary with arousal. Using a previously established sleep scoring procedure (***Novitskaya et al., 2016***; ***Yang et al., 2019***) (Figure 5-figure supplement 1), we classified rat behavior as awake (37.4% ± 2.9% of total recording time), NREM sleep (53.1% ± 3.2%), or REM sleep (1.2% ± 0.9%). Since ripples were extremely rare or essentially absent during REM sleep (not shown), we limited further analysis to ripples occurring during awake (awRipple) or NREM sleep (NREM-Ripple). Expectedly, LC activity was overall higher during awake than during NREM sleep (Figure 5A). The LC suppression occurred around both awRipples and NREM-Ripples, although the dynamic range was bigger during wakefulness. Moreover, during awRipples, LC activity did not decrease to the levels observed during NREM sleep (Figure 5A). Furthermore, peri-ripple LC modulation occurred at slower and faster temporal scales, regardless of the behavioral state. Figure 5B illustrates a slow downward drift in the LC firing rate preceding ripples, likely reflecting the ongoing brain state, as described in the previous Results section. In contrast, event-specific LC modulation had faster dynamics (Figure 5B, highlighted interval). To minimize the influence of global state fluctuations and emphasize event-related dynamics, we applied state correction and z-score normalization for further analysis.

**Figure 5.**
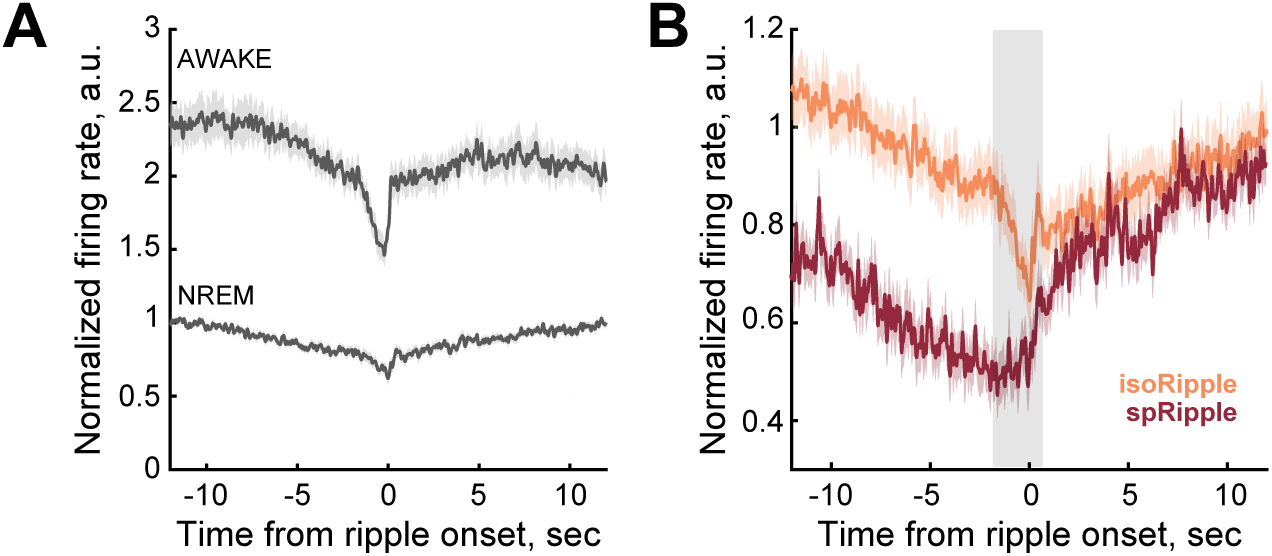
LC modulation around sleep oscillations. **(A)** Peri-ripple LC-MUA during awake state and NREM sleep. LC activity and the range of peri-event LC modulation differed across behavioral states; it was overall higher preceding ripples occurring in wakefulness than in NREM sleep. Despite the state-dependent differences in the firing rate, peri-ripple LC modulation was observed in each behavioral state. During wakefulness, LC activity did not decrease to the levels observed during NREM sleep. **(B)** Peri-event LC-MUA around isolated and spindle-coupled ripples during NREM sleep. LC activity exhibited fast peri-ripple dynamics (highlighted interval) superimposed on slower, state-dependent fluctuations around isolated ripples. Fast LC modulation was absent, while slow fluctuations were preserved around ripples coupled with sleep spindles. For all plots, LC-MUA firing rate was scaled to a pre-event baseline interval [−12 to −10 sec] to preserve baseline differences in LC activity across behavioral states. Bin size: 50 ms. isoRipple – isolated ripple, spRipple - spindle-coupled ripple.

For each data set and each behavioral state, we extracted the percent of ripple subsets with significant LC modulation (subMI >95% CI). We then correlated the subMIs with the total time spent in each behavioral state. The time spent awake, but not in NREM sleep, was significantly correlated with the proportion of ripples associated with LC suppression (r = 0.48, p = 0.03), indicating a more consistent LC modulation around awRipples.

We also examined whether differential peri-ripple LC profiles are related to ripple-spindle temporal coupling, which is known to occur during NREM sleep (***Molle et al., 2006***). We detected sleep spindles (504.5 ± 57.5 per session) occurring at a rate of 8.4 ± 0.8 spindles per minute and split NREM-Ripples into spindle-coupled (spRipple) and isolated (isoRipple). Next, we compared the modulation of the LC around three subtypes of ripples: awRipple (30.1% ± 3.6% of all detected ripples), spRipple (17.2% ± 1.9%), and isoRipple (48.6% ± 3.4%). The ripple subtypes differed in the intra-ripple frequency (Friedman test, *χ*^2^ = 35.62, p < 0.0001, post hoc pairwise comparisons were performed using Wilcoxon signed-rank tests with Holm–Bonferroni correction for multiple comparisons. awRipple vs isoRipple: p = 0.00003 awRipple vs spRipple: p = 0.00004 isoRipple vs spRipple: p = 0.0002), with awRipples being the fastest and spRipples the slowest (Figure 6A). There was no difference in the ripple peak amplitude (Friedman test, *χ*^2^ = 3.7, p = 0.16; Figure 6B). Comparison of the peri-ripple LC MUA rates across three ripple subtypes revealed a significant difference in the proportion of cases showing LC modulation (MI > 95%CI). Specifically, a significant decrease in the LC MUA was detected around awRipples in all 20 sessions (Figure 6C, left), whereas around isoRipples and spRipples, a significant change in LC-MUA was present in 13 (65%) and 4 (20%) sessions, respectively (Figure 6C). This observation was further supported by higher MI values for awRipples, intermediate MIs for isoRipples, and lowest MIs for spRipples (Figure 6D).

**Figure 6.**
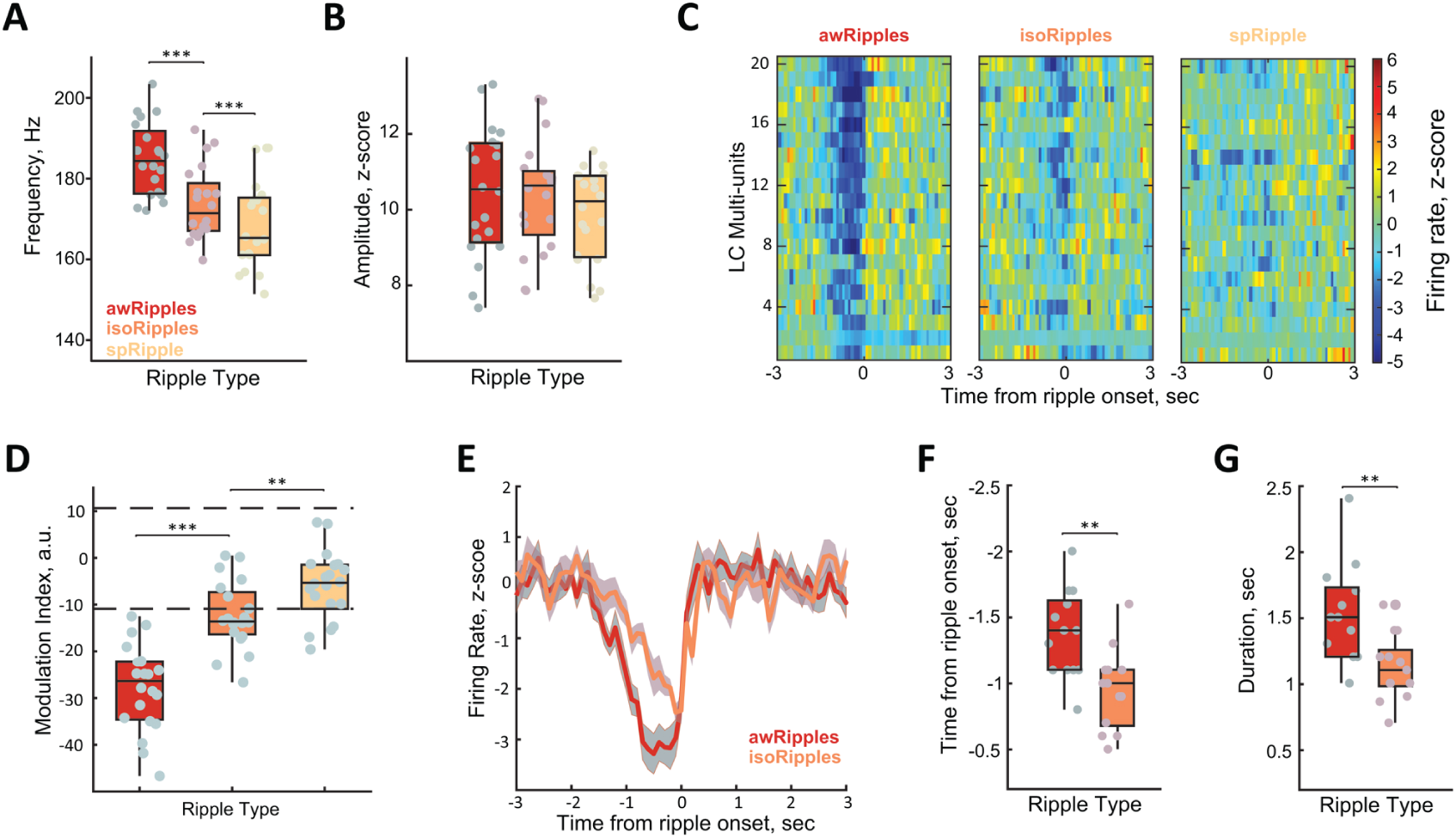
State-dependent modulation of LC activity across ripple subtypes. **(A, B)** Intra-ripple frequency **(A)** and peak amplitude **(B)** for different ripple types. Box-whisker plots show the median, the 1st and 3rd quartiles, and min/max. Gray dots show data from individual rats. *** - p < 0.001 for post hoc pairwise comparisons (Wilcoxon signed-rank tests with Holm–Bonferroni correction for multiple comparisons). **(C, D)** Differential LC activity modulation across ripple types. Session-averaged LC multi-unit activity (MUA) aligned to the ripple onset **(C)** and MI **(D)** is shown for different ripple types. The LC MUA rate is color-coded and plotted for individual sessions. Note the strongest LC activity suppression around ripples occurring in wakefulness (awRipples) and the weakest around ripples coupled with sleep spindles (spRipples). **(E-G)** The patterns of LC activity around different ripple types. The temporal dynamics **(E)**, the onset **(F)**, and the duration **(G)** of ripple-associated LC MUA modulation are shown for ripples occurring during awake (awRipples) and NREM sleep (isoRipples). The data from sessions with significant LC MUA rate decreases are shown. Note an earlier onset and longer duration of LC activity modulation around awake ripples. Due to an overall weak or absent LC activity modulation, data from spindle-coupled ripples are not shown.

Due to the overall very weak or absent LC modulation around spRipples, we compared the LC MUA dynamics around awRipples and isoRipples and only in the sessions showing MI > 95% CI (Figure 6E). This analysis revealed that the onset of the LC spiking suppression occurred significantly earlier preceding awRipples (Wilcoxon signed-rank test, Z = -2.83, p = 0.005; Figure 6F) and the LC suppression was more sustained (Z = -2.80, p = 0.005; Figure 6G).

Thus, the peri-ripple LC modulation depended on the cortical-hippocampal functional connectivity, with LC activity being preserved during the periods of hippocampal-cortical communication, as indicated by the ripple-spindle coupling.

### LC activity modulation around sleep spindles

Consistent with previous studies (***Aston-Jones and Bloom, 1981***; ***Swift et al., 2018***), we observed LC activity modulation around sleep spindles. As expected, both spindle and LC activity were cortical state-dependent (Figure 7A). The correlation analysis confirmed that a higher spindle rate occurred during the EEG epochs with higher SI r = 0.28 to 0.70, p < 0.0001 for all sessions with highest p = 0.00005), Figure 7B, whereas LC-MUA and SI were anticorrelated (r = -0.56 to -0.21, p < 0.01 for all sessions with the highest p = 0.005, Figure 7B). LC-MUA around sleep spindles exhibited bidirectional modulation, characterized by a gradual decrease before spindle onset, followed by a rapid overshoot (Figure 7C, gray line). Similar to peri-ripple LC suppression, a relatively slow (multi-second) temporal profile likely reflected a shift in the thalamo-cortical network preceding sleep spindles. We generated shuffled time series and did not observe any significant LC modulation (not shown). To isolate the LC activity pattern that is not state-dependent, we subtracted LC-MUA aligned to shuffled time series from LC-MUA aligned to spindle onsets. This procedure refined the time window of a transient LC-MUA change (Figure 7C, brown line). Specifically, LC-MUA exhibited a sharp decrease 2.5 ± 0.09 s before spindle onset, followed by a rapid return to baseline and a small overshoot roughly corresponding to the duration of a sleep spindle (1.1 to 2.8 s). Overall, a significant LC modulation (MI > 95% CI) around sleep spindles was present in 17 out of 20 sessions (Figure 7D).

**Figure 7.**
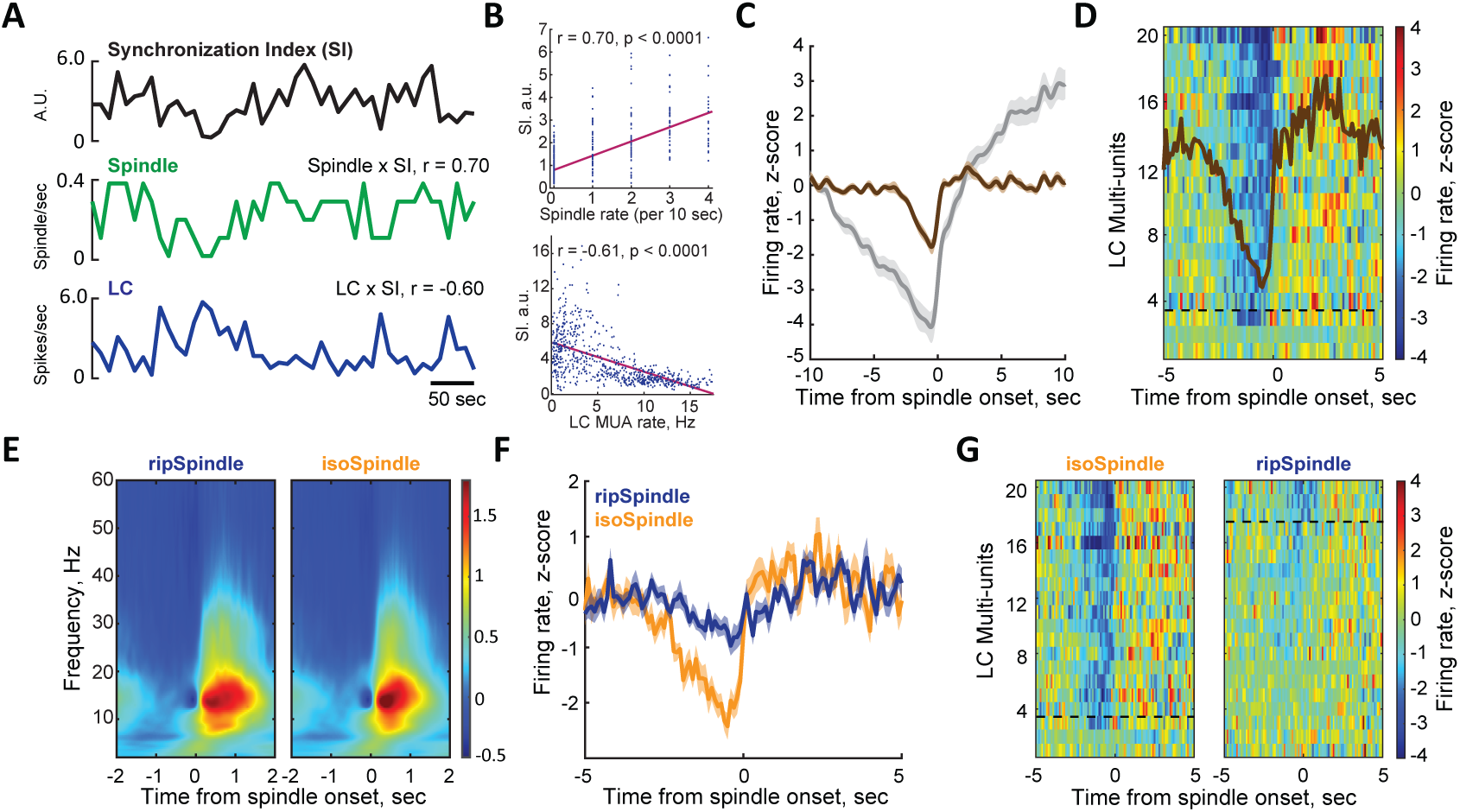
LC activity modulation around sleep spindles. **(A)** Representative traces illustrating the relationship between cortical state, assessed by the Synchronization Index (SI), LC multiunit activity (LC-MUA), and sleep spindle rate. **(B)** Correlation of LC-MUA rate with SI (upper panel) and spindle rate (lower panel) from a representative session. **(C–D)** Spindle-associated LC modulation across slower and faster temporal scales. **(C)** A decrease in LC firing rate was observed as early as 10 s before the spindle onset (gray line). This slow LC modulation was likely driven by fluctuations in ongoing cortical state. To reduce the state-dependent effect, LC-MUA aligned to shuffled spindle times was subtracted. The resulting state-corrected LC-MUA trace (brown) revealed faster LC dynamics. LC-MUA was averaged and smoothed using a 1 Hz low-pass filter. **(D)** Color-coded LC-MUA (state-corrected and z-scored) aligned to the spindle onset for each dataset, with the overlay representing the grand average across sessions (n = 20). **(E)** EEG spectrograms around ripple-coupled (ripSpindle) and isolated (isoSpindle) sleep spindles. The ripSpindle was detected if at least one ripple occurred between the spindle on/offset. Spindle duration was extracted as the time between the spindle on/offset. The ripSpindles were significantly longer (Wilcoxon signed-rank test, p < 0.0001, see Results for more details). **(F–G)** LC activity profiles around ripSpindles and isoSpindles, shown as the grand average **(F)** and session-averaged responses **(G)**. Dashed lines in **(D)** and **(G)** separate cases with significant (top) and non-significant LC modulation.

We next explored if the temporal profile of spindle-associated LC-MUA differed around ripple-coupled (ripSpindle, 41.3 ± 3.1% of all detected spindles) and ripple-uncoupled (isoSpindle, 58.7 ± 3.1%) spindles. Notably, isoSpindles were significantly shorter compared to ripSpindles (1.32 ± 0.05 sec and 1.51 ± 0.06 sec, respectively; Z = –3.92, p = 0.000089; Figure 7E), but two subtypes of sleep spindles did not differ in their oscillatory frequency (ripSpindle: 15.37 ± 0.11 Hz, isoSpindle: 15.33 ± 0.09 Hz; Wilcoxon signed-rank test, Z = –0.22, p = 0.82) or maximum power (ripSpindle: 1.98 ± 0.14 a.u., isoSpindle: 2.08 ± 0.15 a.u.; Z = 1.27, p = 0.20). Despite visible differences in the dynamics of z-scored LC MUA around two types of spindles (Figure 7F), the absolute LC-MUA rate, nor SI differed during a 2-sec time window preceding either group of spindles (Kolmogorov-Smirnov test, p > 0.05 for both variables and all sessions, not shown). Across sessions, MI values exceeded 95% CI in 17/20 datasets for isoSpindles and only 3/20 for ripSpindles. Thus, LC modulation was more consistent and pronounced around isoSpindles compared to ripSpindles (Figures 7F and 7G). Together, this result provided further evidence that LC activity is preserved during hippocampal–cortical communication, as reflected by ripple–spindle coupling.

Lastly, the detection of coupled events (spRipple and ripSpindle) was performed independently, although some overlap cannot be excluded. Figure 7-figure supplement 1 shows a significant correlation (Pearson r = 0.72, p = 0.0003) between the MI around spRipples and ripSpindles. Overall, we found that LC suppression was generally weak around both types of coupled events. Specifically, session-averaged spRipple-associated LC suppression reached a significance level (exceeding 95% CI) in 4 (n = 3 rats) out of 20 sessions. The significant ripSpindle-associated LC suppression was observed in 3 (n = 2 animals) out of 20 sessions.

## Discussion

Using multi-site electrophysiology in freely behaving adult male rats, we characterized the dynamics of LC neuronal activity and hippocampal ripples across two temporal scales. Both LC firing and ripple occurrence were strongly state-dependent, yet inversely related: periods of heightened arousal were marked by increased LC activity and reduced ripple rates. At a finer temporal resolution, LC spiking consistently decreased approximately 1–2 seconds before ripple onset. The magnitude of peri-ripple LC modulation varied across ripple subsets but was not correlated with ripple intrinsic properties. Notably, the strongest LC modulation occurred around ripples in the awake state, whereas LC activity remained largely unchanged around ripples coupled with sleep spindles during NREM sleep. Together, these findings provide novel insight into the state-dependent dynamics of cross-regional interactions and suggest that the LC is involved in a large-scale cortical–subcortical network supporting offline information processing.

### LC-NE dynamics during NREM sleep and its functional implications

LC activity has long been established to fluctuate with arousal level; it is higher during vigilant states and lower during sleep (***Aston-Jones and Bloom, 1981***). This pioneering work has promoted extensive research that has established the critical engagement of the LC-NE system in many cognitive functions dependent on real-time processing of incoming information (***Berridge and Waterhouse, 2003***; ***Sara and Bouret, 2012***; ***Sara, 2009***). However, only recently has the field begun to uncover the temporal dynamics of LC-NE activity and its functional significance during states of low vigilance, such as NREM sleep. Using opto- and chemogenetic tools, it has been confirmed that LC activation induces awakening, whereas LC inhibition promotes sleep (***Carter et al., 2010***; ***Vazey and Aston-Jones, 2014***). Consistent with earlier studies, we observed cortical state-dependent fluctuation of LC neuronal activity in naturally behaving rats. By quantifying cortical state dynamics during NREM sleep at a 4-s resolution, we found an inverse correlation between the LC firing rate and the degree of cortical arousal as indicated by SI. Furthermore, LC activity during NREM sleep was inversely correlated with the incidence of hippocampal ripples and sleep spindles; the latter result is in agreement with inverse relationships between NE release and the power of spindle oscillations (***Osorio-Forero et al., 2021***). Collectively, these observations suggest the existence of a large-scale coordinated network regulating the NREM sleep microstructure and possibly mediating offline information processing.

In our earlier study, by recording LC spiking activity in naturally behaving rats, we observed a delayed time window of enhanced LC neuron firing, specifically during NREM sleep episodes after learning (***Eschenko and Sara, 2008***). Recent studies in mice uncovered infra-slow (∼0.02 Hz) fluctuations of cortical and thalamic NE release during NREM sleep and related the LC activity dynamics to the regulation of sleep microstructure and state transitions (***Osorio-Forero et al., 2021***; ***Kjaerby et al., 2022***). Furthermore, optogenetic LC stimulation at 2Hz during NREM sleep after exposure to a spatial memory task, whilst not affecting ripple occurrence, decreased the stability of replay of hippocampal place cells and caused a memory deficit (***Swift et al., 2018***). This finding aligns with our previous results demonstrating that phasic LC activation time-locked to ripples during post-learning sleep disrupts spatial memory consolidation (***Novitskaya et al., 2016***).

While the mechanisms and functional significance of LC activity patterns during NREM sleep remain to be fully understood (***Foustoukos and Lüthi, 2025***; ***Sara, 2017***; ***Poe, 2017***), these observations align with pharmacological evidence showing that experimental manipulation of NE transmission following learning can alter memory strength (***Przybyslawski et al., 1999***; ***Sara et al., 1999***; ***Roullet and Sara, 1998***; ***Miranda et al., 2009***; ***Gibbs et al., 2010***; ***Gazarini et al., 2013***; ***Gais et al., 2011***). Our recent work suggested a potential network mechanism by which the LC–NE system contributes to sleep-dependent memory consolidation (***Duran et al., 2023***). Specifically, we demonstrated that both *α*2- and *β*-adrenoceptors are involved in the generation of hippocampal ripples. Moreover, pharmacological manipulation of noradrenergic transmission after learning altered all major NREM sleep oscillations and their coupling, potentially leading to less efficient spatial memory consolidation (***Duran et al., 2023***). The present findings extend this work by providing new insights into how LC neuronal activity contributes to the expression of normal sleep microarchitecture.

A more comprehensive investigation of cross-regional coupling and its modulation by the LC–NE system represents an important and still insufficiently explored area of research. Further elucidating the complexity of these interactions will be essential for understanding how noradrenergic signaling shapes large-scale brain network dynamics across behavioral states.

### Coerulear-hippocampal interactions for memory consolidation

Hippocampal ripples are broadly recognized as a central mechanism of memory consolidation (***Klinzing et al., 2019***; ***Skelin et al., 2018***). Emerging work has begun to reveal the circuit mechanisms by which hippocampal ripples engage a larger brain network that includes the prefrontal cortex, thalamus, amygdala, and ventral tegmental area (VTA) for offline information processing (***Girardeau et al., 2017***; ***Latchoumane et al., 2017***; ***Gomperts et al., 2015***; ***Wang and Ikemoto, 2016***; ***Yang et al., 2019***). Our findings expand this memory-supporting circuit to involve the LC-NE system. By characterizing LC and hippocampal activity at a fine temporal scale, we revealed differential patterns of coerulear-hippocampal interactions. First, we observed consistent transient suppression of LC activity occurring 1-2 seconds before the ripple onset. This cessation of LC neuron spiking and the corresponding depletion of NE might have enabled a network to shift to a transient state of enhanced synchronization, allowing ripple generation. This view is indirectly supported by an increase in the EEG delta power immediately preceding the ripple onset, when LC activity is suppressed.

Second, we showed that the degree of LC modulation varied across different subsets of ripples. The strongest and most consistent peri-ripple LC inhibition was observed during the awake state. Interestingly, ripples occurring during wakefulness have been associated with stronger and more structured neuron ensemble reactivation, which was correlated with recent experience (***Tang et al., 2017***). Therefore, a transient silence of LC might be beneficial for the ripple-associated reactivation of recent memory traces. Similarly, coordinated activity of reward-related VTA neurons was greater around ripples that were associated with reactivation of neuronal ensembles that relayed recent learning experience (***Gomperts et al., 2015***). Conversely, LC activation induces arousal, shifts the network to a state incompatible with ripple generation, and thereby might cause interference for the consolidation of recently acquired information (***Novitskaya et al., 2016***; ***Swift et al., 2018***).

During NREM sleep, despite substantially reduced LC activity, we observed a fine-tuned LC activity dynamics around ripples. Specifically, LC neuron discharge was only mildly reduced around isolated ripples, and it was largely preserved during ripples co-occurring with sleep spindles. Differential firing patterns for spindle-coupled and -uncoupled ripples have been reported for cortical and thalamic neurons (***Peyrache et al., 2011***; ***Varela et al., 2001***; ***Yang et al., 2019***). During NREM sleep, hippocampal ripples occur in coordination with cortical slow oscillations and sleep spindles (***Maingret et al., 2016***). The precise temporal correlation between the oscillatory patterns expressed during NREM sleep has been causally linked to memory consolidation by mediating the transfer of newly encoded information from the hippocampus to the cortex for long-term storage (***Latchoumane et al., 2017***; ***Buzsaki, 1996***; ***Molle et al., 2006***; ***Maingret et al., 2016***; ***Klinzing et al., 2019***).

At first glance, the suppression of LC activity around ripples that we observed in the present study may seem inconsistent with the established role of NE in facilitating ripple-mediated synaptic plasticity (***Norimoto et al., 2018***; ***Sadowski et al., 2016***). Remarkably, LC activity was indeed preserved around some subsets of ripples. In our previous work, we showed that LC activation during the time windows marking hippocampal-cortical communication interferes with the cortical dynamics and is detrimental for spatial memory (***Novitskaya et al., 2016***).

Differential modulation of LC activity that was observed in the present study supports the idea that reduced NE release in the hippocampus might bias the content of memory trace during the subsequent ripples and weaken the reactivation of irrelevant experiences. In general, our results highlight the importance of preserved NE transmission at times of cross-regional information transfer. An overall higher LC firing rate during states of elevated arousal may require a larger dynamic range of LC activity modulation (***Kjaerby et al., 2022***). Therefore, in the awake state, a more pronounced suppression of LC activity may be required for the network shift permitting ripple generation.

In summary, our results demonstrate a state-dependent noradrenergic influence on the thalamocortical–hippocampal circuit and position the LC as part of an extended brain network whose coordinated activity may support system-level memory consolidation. Conducting behavioral assays before electrophysiological recordings, along with spatially and temporally precise modulation of LC activity during recording sessions, will be essential for achieving a mechanistic understanding of network dynamics and its functional role for memory consolidation in future investigations. Our present results suggest that during wakefulness, transient suppression of noradrenergic transmission may be associated with the generation of hippocampal ripples and sleep spindles by enhancing network synchrony. We propose that reduced peri-ripple LC activity during the awake state may preferentially promote the replay of recent memory traces by limiting interference from remote memories, which are more likely to be reactivated in more alert states. In contrast, during NREM sleep, preserved LC activity during ripples coinciding with sleep spindles suggests a role for NE in facilitating cross-regional communication underlying memory-related information transfer. These results provide new insight into how the LC–NE system may shape memory processing across brain states and open the way for future causal investigations using modern circuit-level tools.

## Methods and Materials

### Animals

Seven adult male Sprague-Dawley rats (RRID:RGD70508, 350-400 g) were used. Data from one rat has been collected for a previous study **Eschenko et al.** (**2012**) and reanalyzed. The other six rats (Charles River Laboratory, Germany) were kept on a 12-hour light-dark cycle (8:00 am lights on) and single-housed after surgery. Recordings were performed during the dark cycle. Similar surgery and recording procedures were used as reported in detail elsewhere (***Yang et al., 2019***; ***Eschenko et al., 2012***). The study was performed in accordance with the German Animal Welfare Act (TierSchG), Animal Welfare Laboratory Animal Ordinance (TierSchVersV), and ARRIVE guidelines. This is in full compliance with the guidelines of the EU Directive on the protection of animals used for scientific purposes (2010/63/EU). The study was reviewed by the ethics commission (§15 Tier-SchG) and approved by the state authority (Regierungspräsidium, Tübingen, Baden-Württemberg, Germany, permission KY01/19).

### Surgery and electrode placement

Procedures for stereotaxic surgery under isoflurane anesthesia have been described in detail elsewhere (***Yang et al., 2019***). Briefly, anesthesia was initiated with 4% and maintained with 1.5-2.0% isoflurane. Body temperature, heart rate, and blood oxygenation were monitored throughout the entire anesthesia period. The depth of anesthesia was controlled by a lack of pain and sensory response (hind paw pinch). A fully anesthetized rat was fixed in a stereotaxic frame with the head angle at zero degrees. The skull was exposed and a local anesthetic (Lidocard 2%, B. Braun, Melsungen, Germany) was applied to the skin edges. Craniotomies were performed on the right hemisphere above the target regions. Additional burr holes were made for EEG, grown, and anchor screws (stainless steel, 0.86-1.19 mm diameter, Fine Science Tools, Heidelberg, Germany). Dura mater was removed when necessary. For extracellular recording, single platinum-iridium electrodes (FHC, Bowdoin, ME) were placed in the anterior cingulate cortex (ACC, AP/ML: 2.8 mm/0.8mm from Bregma and DV: 1.8 mm from the dura surface). For the recording of the hippocampal ripples, the electrode was mounted on a self-made movable microdrive and inserted above the CA1 subfield of the dorsal hippocampus (dHPC, -3mm/2mm/ 2mm). For LC recordings, single platinum–iridium electrodes (FHC, Bowdoin, ME; n = 1), microwire brush electrodes (Micro-Probes, MD; n = 2), and silicon probes (Cambridge Neurotech, Cambridge, UK; n = 2) were mounted on a microdrive (Cambridge Neurotech, Cambridge, UK) and implanted above the LC (4.2 mm posterior, 1.2 mm lateral from lambda; DV: 5.5–6.2 mm) at a 15° angle. We also implanted one rat with a polymer electrode array (LLNL, Livermore, CA). The accuracy of LC targeting was verified by online monitoring of neural activity. The LC neurons were identified by broad spike widths (0.6 msec), regular low firing rate (1 –2 spikes/s), and a biphasic (excitation followed by inhibition) response to paw pinch (***Eschenko et al., 2012***). For EEG recording, a screw was placed above the frontal cortex. The ground screw was placed above the cerebellum. Screws were fixed in the skull and additionally secured with tissue adhesive. The entire implant was secured on the skull with dental cement (RelyX™ Unicem 2 Automix, 3M, MN). A copper mesh was mounted around the implant to shield and protect the exposed connection wires. During the post-surgery recovery period, analgesic (2.5 mg/kg, s.c.; Rimadyl, Zoetis) and antibiotic (5.0 mg/kg, s.c.; Baytril®) were given for 3 and 5 days, respectively. Electrode placements were histologically verified.

### Electrophysiological recording and data analysis

Rats were first habituated to the sleeping box (30 x 30 x 40 cm) and the cable plugging procedure. The electrodes were either connected to the Neuralynx Digital Lynx or FreeLynx wireless acquisition system via two 32-channel head stages (Neuralynx, Bozeman, MT). The electrode placement in the LC was optimized by lowering the electrodes with a maximal 0.05 mm step and monitoring spiking activity on the high-passed (300 Hz – 8kHz) extracellular signal. The depth of the dHPC electrodes was adjusted by gradually lowering the electrode (maximum 0.05 mm per day) until reliable ripple activity was observed. Once the electrode position was optimized, the broadband (0.1Hz - 8kHz) extracellular signals were acquired and digitized at 32kHz and referenced to the ground screw. The animals’ movement was monitored by an EMG electrode attached to the neck muscle. All recordings were performed between 10 a.m. and 7 p.m.

### Classification of behavioral states

We classified the rat’s spontaneous behavior into awake and NREM sleep using frontal EEG or cortical LFPs by applying a previously established sleep scoring algorithm (***Novitskaya et al., 2016***; ***Yang et al., 2019***). Briefly, animal movement speed was extracted from the video recording synchronized with a neural signal. Artefact-free EEG signals were band-pass filtered using a Butterworth filter implemented in Matlab 2024a (MathWorks, Natick, MA). Subsequently, delta-band power (*δ*, 1–4 Hz), theta-band power (*θ*, 6–10 Hz), and the *θ∕δ* power ratio were computed within contiguous 4-second epochs. The epochs of the awake state were identified by the presence of active locomotion and *θ∕δ* values above the arbitrarily-defined threshold; the epochs of NREM sleep were identified by the absence of motor activity and *θ∕δ* values below the threshold. The minimal duration of the same behavioral state was set to 20 sec; data segments with less steady behavioral states were excluded from further analysis.

### Event detection

For detection of the hippocampal ripples, a broadband (0.1 Hz - 8 kHz) extracellular signal recorded from the dHPC (pyramidal layer, CA1 subfield) was band-pass (120 – 250 Hz) filtered, rectified, and low-pass filtered (25 Hz). The resulting signal was z-score normalized, and ripple oscillations were detected by signal thresholding at 5 standard deviations (SDs). Ascending and descending crossings at 1 SD defined the ripple onset and offset, respectively. Clustered and isolated ripples were classified based on the inter-ripple interval (IRI) as described in detail elsewhere (***Selinger et al., 2007***). Briefly, IRIs were extracted, and the log(IRI) distribution was analyzed. A bimodal log(IRI) distribution indicated ripple occurrence with shorter and longer IRI. A crossing point of two distributions was used for classifying the ripple type. Ripples with short IRI (< 0.33 sec) were classified as ‘clustered’ and treated as a single ripple event; the onset time was defined by the first ripple in a cluster. Ripples with IRI > 0.33 sec were classified as ‘isolated’.

For the detection of sleep spindles, the EEG signal was band-pass (12 - 16 Hz) filtered, down-sampled (200 Hz), and the root mean square (0.2 s smoothing) was calculated. The spindle detection threshold was defined at 3 SD of the signal amplitude during NREM sleep episodes. Ascending and descending signal crossings at 1SD defined the spindle on- and offset, respectively. The spindle duration was extracted as the time between the spindle onset and offset; the minimal sleep spindle duration was set to 0.5 sec. Coupled oscillatory events were defined as ripples occurring between spindle onset and offset.

### Analysis of peri-event neural activity

For analysis of LC activity modulation around ripples, we used a previously established method (***Yang et al., 2019***). First, the spike times of LC multiunit activity (LC-MUA) were extracted by high-pass (600Hz) filtering of the broadband extracellular signal recorded from LC and thresholding at -0.05 mV. The LC-MUA was triggered on ripple/spindle onset, and the averaged peri-event spike histogram (PETH) was generated either ± 6 sec or ± 3 sec around all detected ripples and ± 5 sec around sleep spindles (5-ms bins, Gaussian smoothed with a 10-ms window). To compensate for possible state-dependent fluctuations of LC-MUA, a ‘surrogate’ event sequence was generated by randomly distributing the same number of time points within a 4-second time window around each event; the jittering procedure was repeated 100 times. The averaged PETH was generated around jittered events. Time series of LC MUA around the jittered events were subtracted from the corresponding values around real events, and the resulting (‘corrected’) PETHs were z-score normalized. To further smooth the PETH, we extracted the area above/below the curve of each PETH every 100 ms and z-scored the modulation response. To quantify peri-event dynamics of LC-MUA, a modulation index (MI) was calculated by modulation response within 1 sec before the event onsets. To determine the significance of modulation, we calculated the modulation responses around ‘shuffled’ ripple times generated by permutation of the IRIs and calculated the MIs for each of the 5000 shuffled PETHs; a 95% confidence interval (CI) served as the significance threshold. To quantify the LC-MUA modulation around subsets of ripples, subsampled MI (subMI) was computed for a subset of ripples (20% of all detected ripples in each session), and the same procedure was repeated 5000 times. The s-MI distribution was analyzed.

### Spectral analysis

For LFP analysis, we applied the Morlet-wavelet time-frequency analysis to estimate spectral power around ripple onsets. To compare delta and spindle band activity preceding ripples associated with either strong or weak LC activity modulation, we first averaged the spectral power within the delta (1–4 Hz) and spindle (12–16 Hz) frequency bands over a 2-s window before ripple onset. We then quantified the power by calculating the area under the curve for each frequency band.

### Statistical Analysis

All statistical analyses were performed in MATLAB (RRID:SCR001622, MathWorks, Natick, MA). The analysis of variance (one-way ANOVA) or the Kolmogorov–Smirnov test was used for between-group comparisons, with significance defined as *p* < 0.05. The Wilcoxon signed-rank test was used to compare two matched samples. The Friedman test was used to compare intrinsic ripple properties. Post hoc analyses were performed using the Wilcoxon signed-rank test with Holm–Bonferroni correction for multiple comparisons (*p* < 0.05 after correction). Pearson correlation was used to assess relationships between variables, with significance defined as *p* < 0.05.

## Data Availability

The electrophysiological data and MATLAB code used to generate the results reported in this study are available from the corresponding author upon reasonable request.

## Acknowledgments

This work was funded by the Max Planck Society. We thank Nikos Logothetis for his support of this research project, Susan Sara for her scientific inspiration and insightful discussions, and Loren Frank for providing polymer electrode arrays and valuable advice regarding their implantation. We are also grateful to Axel Oeltermann and Joachim Werner for their expert technical assistance.

## Supplementary Materials

**Figure 1-figure supplement 1.**
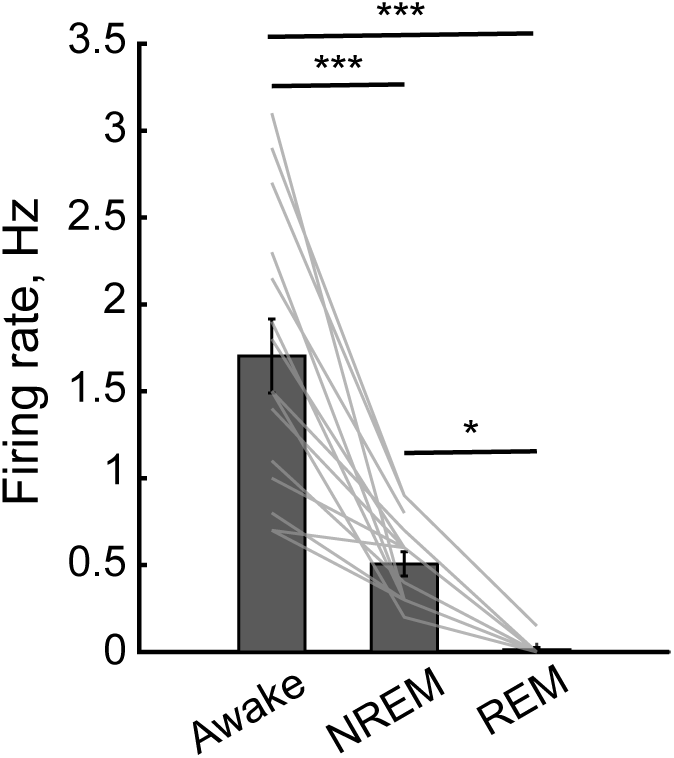
Firing rates of LC neurons across sleep/awake states. Bars show the grand means (± SE) and gray lines show session-averages for each rat. * p < 0.05, ** p < 0.01, and *** p < 0.001 (one-way ANOVA).

**Figure 2-figure supplement 1.**
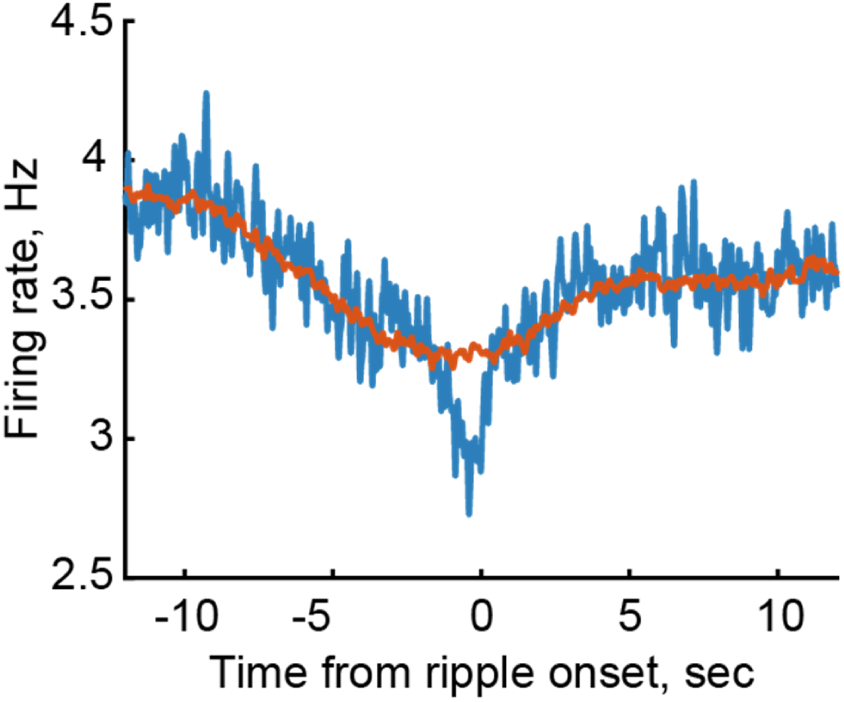
Slow-timescale modulation of ripple-associated LC activity. The LC firing rate (blue trace; bin size: 0.05 s) is plotted ±12 s around ripple onset for a representative session. Notably, a decrease in LC firing rate emerges as early as 10 s prior to ripple onset.

**Figure 3-figure supplement 1.**
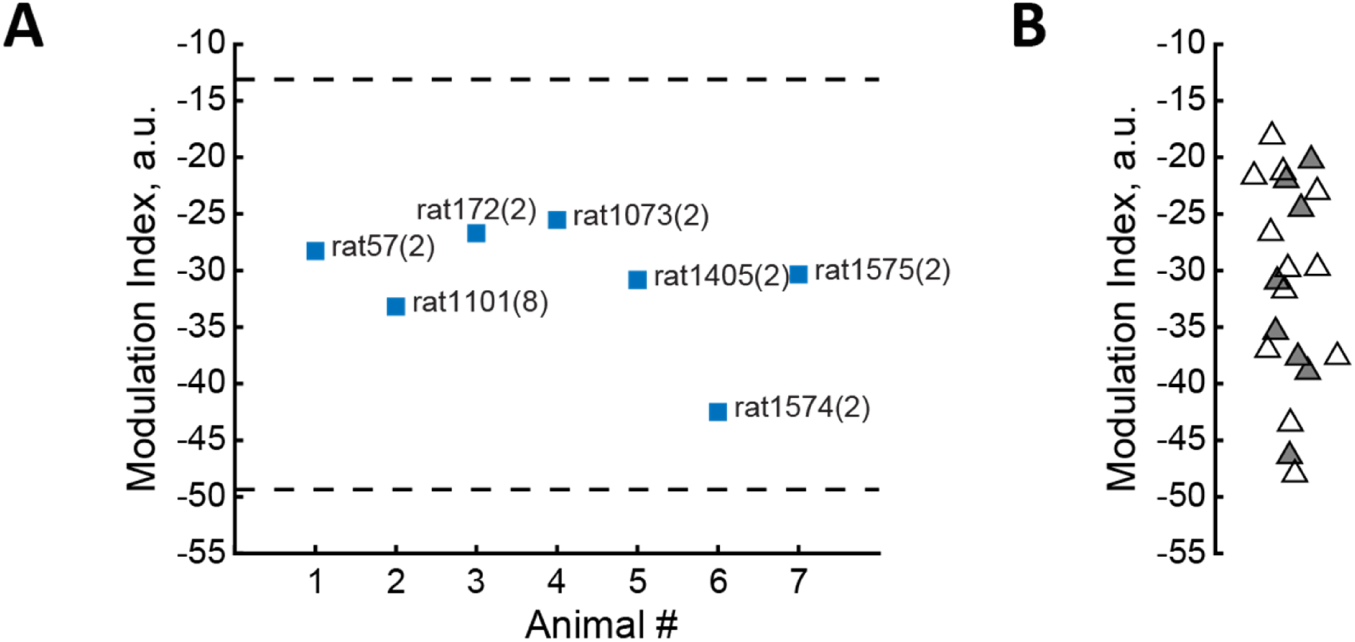
Distribution of the modulation index (MI) across rats and sessions. **(A)** The average MI is shown for each animal, with rat ID and number of sessions indicated in parentheses. Dashed lines denote the mean ± 2 standard deviations across all sessions. Animal-averaged MIs fall within a consistent range, indicating that the distribution is not driven by a single subject (e.g., rat 1101, 8 sessions). **(B)** The average MI is shown for each session. The eight sessions from rat 1101 are indicated by gray-filled triangles.Comparison of these sessions with the remaining 12 sessions from six other animals revealed no significant difference in MI distribution (Kolmogorov–Smirnov test, p = 0.969).

**Figure 5-figure supplement 1.**
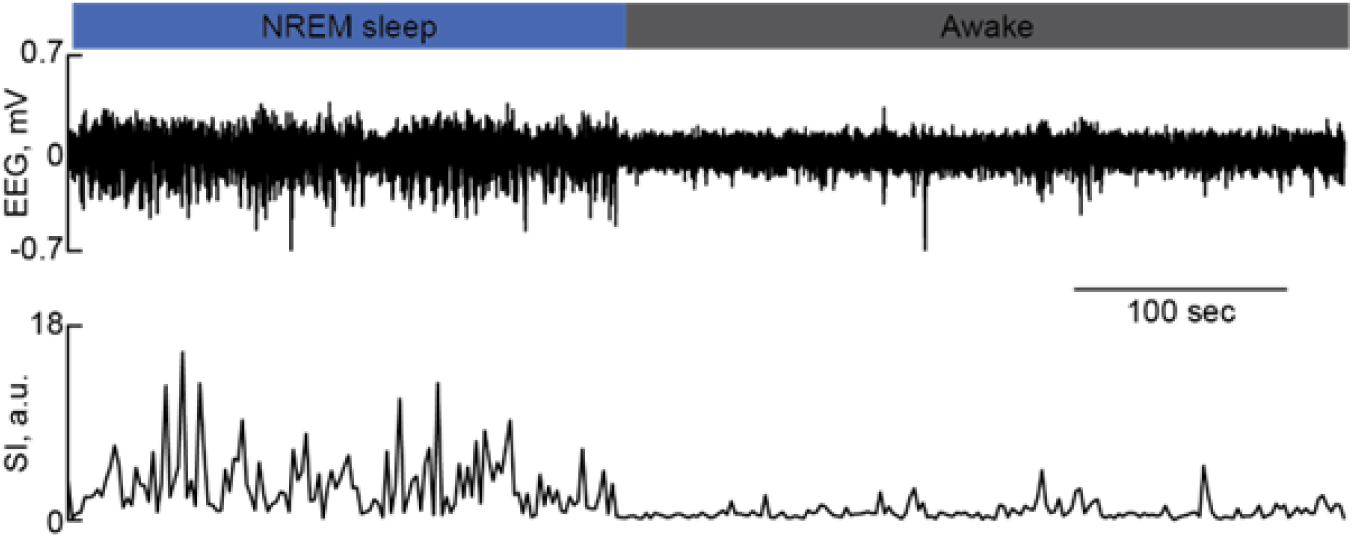
Sleep scoring procedure. Classified NREM sleep and wake episodes (top), with corresponding raw EEG traces (1–300 Hz; middle) and Synchronization Index (SI; bottom), calculated over 4-s epochs.

**Figure 7-figure supplement 1.**
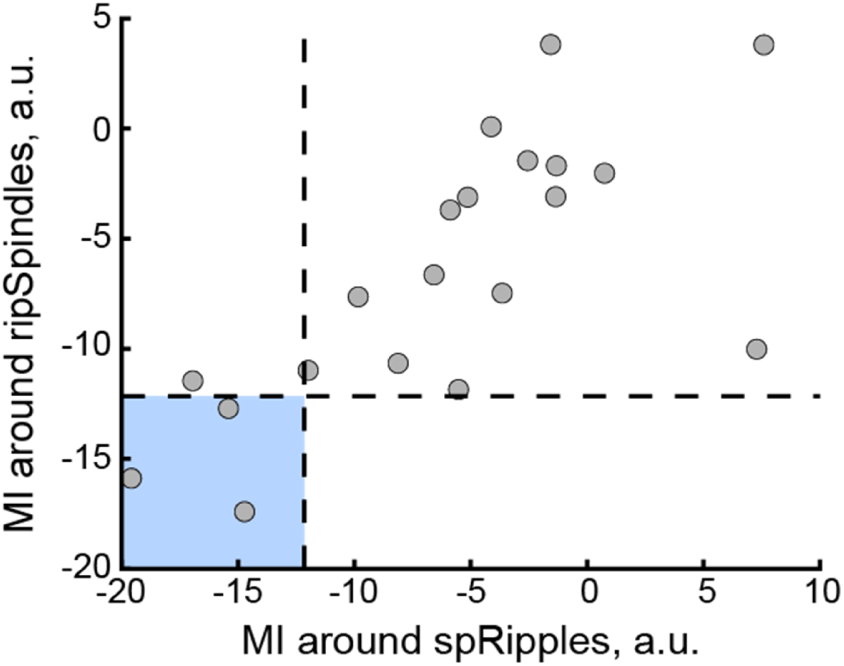
LC modulation around coupled oscillations. Pearson correlation between the modulation index (MI) for spindle-coupled ripples (spRipple) and ripple-coupled spindles (ripSpindle). Detection of coupled oscillations (spRipple and ripSpindle) was performed independently, although some overlap cannot be excluded. Blue square highlights three sessions exhibiting significant (MI < 95% CI) LC suppression around both ripSpindles and spRipples. Generally, LC modulation around coupled oscillations was weak. Specifically, the LC suppression around spRipples and ripSpindles reached significance in 4 sessions (from 3 rats) and 3 sessions (from 2 rats), respectively, out of a total of 20 sessions (from 7 rats).

## References

Aston-Jones G, Bloom FE. Activity of norepinephrine-containing locus coeruleus neurons in behaving rats anticipates fluctuations in the sleep-waking cycle. J Neurosci. 1981; 1(8):876–86. doi: 10.1523/jneurosci.01-08-00876.1981.

Berridge CW, Waterhouse BD. The locus coeruleus-noradrenergic system: modulation of behavioral state and state-dependent cognitive processes. Brain Res Brain Res Rev. 2003; 42(1):33–84. http://www.ncbi.nlm.nih.gov/entrez/query.fcgi?cmd=Retrieve&db=PubMed&dopt=Citation&list_uids=12668290.

Brodt S, Inostroza M, Niethard N, Born J. Sleep-A brain-state serving systems memory consolidation. Neuron. 2023; 111(7):1050–1075. https://linkinghub.elsevier.com/retrieve/pii/S0896627323002015, doi: 10.1016/j.neuron.2023.03.005, clonidine Project.

Buzsaki G. The hippocampo-neocortical dialogue. Cereb Cortex. 1996; 6(2):81–92. http://www.ncbi.nlm.nih.gov/entrez/query.fcgi?cmd=Retrieve&db=PubMed&dopt=Citation&list_uids=8670641.

Cahill L, Prins B, Weber M, McGaugh JL. Beta-adrenergic activation and memory for emotional events. Nature. 1994; 371.

Carter ME, Yizhar O, Chikahisa S, Nguyen H, Adamantidis A, Nishino S, Deisseroth K, de Lecea L. Tuning arousal with optogenetic modulation of locus coeruleus neurons. Nat Neurosci. 2010; 13(12):1526–33. https://www.ncbi.nlm.nih.gov/pubmed/21037585, doi: 10.1038/nn.2682.

Clayton EC, Williams CL. Noradrenergic receptor blockade of the NTS attenuates the mnemonic effects of epinephrine in an appetitive light-dark discrimination learning task. Neurobiol Learn Mem. 2000; 74(2):135–45. http://www.ncbi.nlm.nih.gov/pubmed/10933899, doi: 10.1006/nlme.1999.3946.

Duran E, Pandinelli M, Logothetis NK, Eschenko O. Altered norepinephrine transmission after spatial learning impairs sleep-mediated memory consolidation in rats. Sci Rep. 2023; 13(1):4231. https://www.ncbi.nlm.nih.gov/pubmed/36918712 https://www.ncbi.nlm.nih.gov/pmc/articles/PMC10014950/pdf/41598_2023_Article_31308.pdf, doi: 10.1038/s41598-023-31308-1, duran, Ernesto Pandinelli, Martina Logothetis, Nikos K Eschenko, Oxana eng Research Support, Non-U.S. Gov’t England 2023/03/16 Sci Rep. 2023 Mar 14;13(1):4231. doi: 10.1038/s41598-023-31308-1.

Eschenko O, Magri C, Panzeri S, Sara SJ. Noradrenergic neurons of the locus coeruleus are phase locked to cortical up-down states during sleep. Cereb Cortex. 2012; 22(2):426–35. https://www.ncbi.nlm.nih.gov/pubmed/21670101, doi: 10.1093/cercor/bhr121.

Eschenko O, Sara SJ. Learning-dependent, transient increase of activity in noradrenergic neurons of locus coeruleus during slow wave sleep in the rat: brain stem-cortex interplay for memory consolidation? Cereb Cortex. 2008; 18(11):2596–603. https://www.ncbi.nlm.nih.gov/pubmed/18321875, doi: 10.1093/cercor/bhn020.

Foustoukos G, Lüthi A. Monoaminergic signaling during mammalian NREM sleep - Recent insights and next-level questions. Curr Opin Neurobiol. 2025; 92:103025. doi: 10.1016/j.conb.2025.103025, 1873-6882 Foustoukos, Georgios Lüthi, Anita Journal Article Review England 2025/04/24 Curr Opin Neurobiol. 2025 Jun;92:103025. doi: 10.1016/j.conb.2025.103025. Epub 2025 Apr 22.

Gais S, Rasch B, Dahmen JC, Sara S, Born J. The Memory Function of Noradrenergic Activity in Non-REM Sleep. Journal of Cognitive Neuroscience. 2011; 23(9):2582–2592. https://doi.org/10.1162/jocn.2011.21622, doi: 10.1162/jocn.2011.21622.

Galeotti N, Bartolini A, Ghelardini C. Alpha-2 agonist-induced memory impairment is mediated by the alpha-2A-adrenoceptor subtype. Behavioural Brain Research. 2004; 153(2):409–417. https://www.sciencedirect.com/science/article/pii/S0166432803004856, doi: 10.1016/j.bbr.2003.12.016.

Gazarini L, Stern CAJ, Carobrez AP, Bertoglio LJ. Enhanced noradrenergic activity potentiates fear memory consolidation and reconsolidation by differentially recruiting *α*1- and *β*-adrenergic receptors. Learning & Memory. 2013; 20(4):210–219. http://learnmem.cshlp.org/content/20/4/210.abstract, doi: 10.1101/lm.030007.112.

Gelinas JN, Nguyen PV. *β*-Adrenergic Receptor Activation Facilitates Induction of a Protein Synthesis-Dependent Late Phase of Long-Term Potentiation. The Journal of Neuroscience. 2005; 25(13):3294–3303. http://www.jneurosci.org/content/25/13/3294.abstract http://www.jneurosci.org/content/jneuro/25/13/3294.full.pdf, doi: 10.1523/jneurosci.4175-04.2005.

Gibbs ME, Hutchinson DS, Summers RJ. Noradrenaline release in the locus coeruleus modulates memory formation and consolidation; roles for *α*- and *β*-adrenergic receptors. Neuroscience. 2010; 170(4):1209–1222. https://www.sciencedirect.com/science/article/pii/S0306452210010559, doi: 10.1016/j.neuroscience.2010.07.052.

Girardeau G, Inema I, Buzsaki G. Reactivations of emotional memory in the hippocampus-amygdala system during sleep. Nat Neurosci. 2017; 20(11):1634–1642. http://www.ncbi.nlm.nih.gov/pubmed/28892057 https://www.nature.com/articles/nn.4637.pdf, doi: 10.1038/nn.4637.

Gomperts SN, Kloosterman F, Wilson MA. VTA neurons coordinate with the hippocampal reactivation of spatial experience. Elife. 2015; 4. http://www.ncbi.nlm.nih.gov/pubmed/26465113 https://www.ncbi.nlm.nih.gov/pmc/articles/PMC4695386/pdf/elife-05360.pdf, doi: 10.7554/eLife.05360.

Groch S, Wilhelm I, Diekelmann S, Sayk F, Gais S, Born J. Contribution of norepinephrine to emotional memory consolidation during sleep. Psychoneuroendocrinology. 2011; 36(9):1342–1350. http://www.sciencedirect.com/science/article/pii/S0306453011000953, doi: 10.1016/j.psyneuen.2011.03.006.

Hagena H, Hansen N, Manahan-Vaughan D. *β*-Adrenergic Control of Hippocampal Function: Sub-serving the Choreography of Synaptic Information Storage and Memory. Cerebral Cortex. 2016; http://cercor.oxfordjournals.org/content/early/2016/01/23/cercor.bhv330.abstract http://cercor.oxfordjournals.org/content/26/4/1349.full.pdf, doi: 10.1093/cercor/bhv330.

Hansen N, Manahan-Vaughan D. Locus Coeruleus Stimulation Facilitates Long-Term Depression in the Dentate Gyrus That Requires Activation of *β*-Adrenergic Receptors. Cerebral Cortex. 2015; 25(7):1889–1896. https://doi.org/10.1093/cercor/bht429, doi: 10.1093/cercor/bht429.

Harley CW. Norepinephrine and the dentate gyrus. In: Scharfman HE, editor. The Dentate Gyrus: A Comprehensive Guide to Structure, Function, and Clinical Implications, vol. 163 of Progress in Brain Research Elsevier; 2007.p. 299–318. https://www.sciencedirect.com/science/article/pii/S0079612307630180, doi: 10.1016/S0079-6123(07)63018-0.

Hayat H, Regev N, Matosevich N, Sales A, Paredes-Rodriguez E, Krom AJ, Bergman L, Li Y, Lavigne M, Kremer EJ, Yizhar O, Pickering AE, Nir Y. Locus coeruleus norepinephrine activity mediates sensory-evoked awakenings from sleep. Science Advances. 2020; 6(15):eaaz4232. https://advances.sciencemag.org/lookup/doi/10.1126/sciadv.aaz4232, doi: 10.1126/sciadv.aaz4232, lC and ACC LC PPI Tech paper.

Kjaerby C, Andersen M, Hauglund N, Untiet V, Dall C, Sigurdsson B, Ding F, Feng J, Li Y, Weikop P, Hirase H, Nedergaard M. Memory-enhancing properties of sleep depend on the oscillatory amplitude of norepinephrine. Nature Neuroscience. 2022; 25(8):1059–1070. https://www.nature.com/articles/s41593-022-01102-9, doi: 10.1038/s41593-022-01102-9.

Klinzing JG, Niethard N, Born J. Mechanisms of systems memory consolidation during sleep. Nat Neurosci. 2019; 22(10):1598–1610. https://www.ncbi.nlm.nih.gov/pubmed/31451802, doi: 10.1038/s41593-019-0467-3.

Kuffel A, Eikelmann S, Terfehr K, Mau G, Kuehl LK, Otte C, Löwe B, Spitzer C, Wingenfeld K. Noradrenergic blockade and memory in patients with major depression and healthy participants. Psychoneuroendocrinology. 2014; 40:86–90. https://www.sciencedirect.com/science/article/pii/S0306453013004046, doi: 10.1016/j.psyneuen.2013.11.001.

Latchoumane CV, Ngo HV, Born J, Shin HS. Thalamic Spindles Promote Memory Formation during Sleep through Triple Phase-Locking of Cortical, Thalamic, and Hippocampal Rhythms. Neuron. 2017; 95(2):424–435 e6. http://www.ncbi.nlm.nih.gov/pubmed/28689981 https://ac.els-cdn.com/S0896627317305494/1-s2.0-S0896627317305494-main.pdf?_tid=51e77677-0532-4cd7-be7b-21f7f0e6c4c9&acdnat=1532076239_c347db300f2648a9fef30c6c14c84a51, doi: 10.1016/j.neuron.2017.06.025.

Logothetis NK. Neural-Event-Triggered fMRI of large-scale neural networks. Curr Opin Neurobiol. 2015; 31:214–22. http://www.ncbi.nlm.nih.gov/pubmed/25536423, doi: 10.1016/j.conb.2014.11.009.

Logothetis NK, Eschenko O, Murayama Y, Augath M, Steudel T, Evrard HC, Besserve M, Oeltermann A. Hippocampal-cortical interaction during periods of subcortical silence. Nature. 2012; 491(7425):547–53. https://www.ncbi.nlm.nih.gov/pubmed/23172213, doi: 10.1038/nature11618.

Maingret N, Girardeau G, Todorova R, Goutierre M, Zugaro M. Hippocampo-cortical coupling mediates memory consolidation during sleep. Nat Neurosci. 2016; 19(7):959–964. https://doi.org/10.1038/nn.4304, doi: 10.1038/nn.4304 http://www.nature.com/neuro/journal/v19/n7/abs/nn.4304.html##supplementary-information.

Miranda MI, Ortiz-Godina F, García D. Differential involvement of cholinergic and beta-adrenergic systems during acquisition, consolidation, and retrieval of long-term memory of social and neutral odors. Behavioural Brain Research. 2009; 202(1):19–25. https://www.sciencedirect.com/science/article/pii/S0166432809001545, doi: 10.1016/j.bbr.2009.03.008.

Molle M, Yeshenko O, Marshall L, Sara SJ, Born J. Hippocampal sharp wave-ripples linked to slow oscillations in rat slow-wave sleep. J Neurophysiol. 2006; 96(1):62–70. https://www.ncbi.nlm.nih.gov/pubmed/16611848, doi: 10.1152/jn.00014.2006.

Nitzan N, Swanson R, Schmitz D, Buzsáki G. Brain-wide interactions during hippocampal sharp wave ripples. Proc Natl Acad Sci U S A. 2022; 119(20):e2200931119. doi: 10.1073/pnas.2200931119.

Norimoto H, Makino K, Gao M, Shikano Y, Okamoto K, Ishikawa T, Sasaki T, Hioki H, Fujisawa S, Ikegaya Y. Hippocampal ripples down-regulate synapses. Science. 2018; 359(6383):1524–1527. https://science.sciencemag.org/content/sci/359/6383/1524.full.pdf, doi: 10.1126/science.aao0702.

Novitskaya Y, Sara SJ, Logothetis NK, Eschenko O. Ripple-triggered stimulation of the locus coeruleus during post-learning sleep disrupts ripple/spindle coupling and impairs memory consolidation. Learn Mem. 2016; 23(5):238–48. https://www.ncbi.nlm.nih.gov/pubmed/27084931, doi: 10.1101/lm.040923.115.

Osorio-Forero A, Cardis R, Vantomme G, Guillaume-Gentil A, Katsioudi G, Devenoges C, Fernandez LMJ, Lüthi A. Noradrenergic circuit control of non-REM sleep substates. Current Biology. 2021; 31(22):5009–5023.e7. https://www.sciencedirect.com/science/article/pii/S0960982221012811, doi: 10.1016/j.cub.2021.09.041.

Palacios-Filardo J, Mellor JR. Neuromodulation of hippocampal long-term synaptic plasticity. Current Opinion in Neurobiology. 2019; 54:37–43. http://www.sciencedirect.com/science/article/pii/S0959438818301041, doi: 10.1016/j.conb.2018.08.009.

Paxinos G, Watson C. The rat brain in stereotaxic coordinates. 5th ed. Amsterdam; Boston: Elsevier Academic Press; 2005. http://www.loc.gov/catdir/description/els051/2004303983.html, 2004303983 GBA469799 George Paxinos, Charles Watson. chiefly ill.; 29 cm. + 1 CD-ROM (4 3/4 in.) System requirements for accompanying disc: Windows: Pentium II processor 233 MHz or higher; 64 MB RAM; Windows 98/NT4.0/2000/XP; Adobe Acrobat Reader 5.0 or later; VGA monitor with 256 colors; 800 x 600 screen resolution; 8X or faster CD-ROM drive. System requirements for accompanying disc: Macintosh: Aple PowerPC G3 333 MHz or higher; 128 MB RAM (256 recommended); Mac OS 9.x or higher (OS X recommended); Adobe Acrobat Reader 5.0 or later; VGA monitor with 256 colors; 800 x 600 screen resolution; 8X or faster CD-ROM drive. Includes bibliographical references (p. xxv-xxix) and index.

Peyrache A, Battaglia FP, Destexhe A. Inhibition recruitment in prefrontal cortex during sleep spindles and gating of hippocampal inputs. Proceedings of the National Academy of Sciences. 2011; 108(41):17207–17212. http://www.pnas.org/content/pnas/108/41/17207.full.pdf, doi: 10.1073/pnas.1103612108.

Poe GR. Sleep Is for Forgetting. The Journal of Neuroscience. 2017; 37(3):464–473. doi: 10.1523/jneurosci.0820-16.2017.

Przybyslawski J, Roullet P, Sara SJ. Attenuation of emotional and nonemotional memories after their reactivation: role of beta adrenergic receptors. J Neurosci. 1999; 19(15):6623–8. http://www.ncbi.nlm.nih.gov/entrez/query.fcgi?cmd=Retrieve&db=PubMed&dopt=Citation&list_uids=10414990.

Roullet P, Sara S. Consolidation of memory after its reactivation: involvement of beta noradrenergic receptors in the late phase. Neural Plast. 1998; 6(3):63–8. http://www.ncbi.nlm.nih.gov/entrez/query.fcgi?cmd=Retrieve&db=PubMed&dopt=Citation&list_uids=9920683 https://www.ncbi.nlm.nih.gov/pmc/articles/PMC2565312/pdf/NP-06-03-063.pdf.

Sadowski JLP, Jones M, Mellor J. Sharp-Wave Ripples Orchestrate the Induction of Synaptic Plasticity during Reactivation of Place Cell Firing Patterns in the Hippocampus. Cell Reports. 2016; 14(8):1916–1929. http://www.sciencedirect.com/science/article/pii/S2211124716300390 http://ac.els-cdn.com/S2211124716300390/1-s2.0-S2211124716300390-main.pdf?_tid=e72a8314-85a2-11e6-9127-00000aab0f27&acdnat=1475084719_459bd79e6a1810ef6fff395a2ff41a66, doi: 10.1016/j.celrep.2016.01.061.

Sara SJ. The locus coeruleus and noradrenergic modulation of cognition. Nat Rev Neurosci. 2009; 10(3):211–23. http://www.ncbi.nlm.nih.gov/entrez/query.fcgi?cmd=Retrieve&db=PubMed&dopt=Citation&list_uids=19190638 https://www.nature.com/articles/nrn2573.pdf, doi: nrn2573 [pii] 10.1038/nrn2573.

Sara SJ, Roullet P, Przybyslawski J. Consolidation of memory for odor-reward association: beta-adrenergic receptor involvement in the late phase. Learn Mem. 1999; 6(2):88–96. http://www.ncbi.nlm.nih.gov/entrez/query.fcgi?cmd=Retrieve&db=PubMed&dopt=Citation&list_uids=10327234 https://www.ncbi.nlm.nih.gov/pmc/articles/PMC311281/pdf/x1.pdf.

Sara SJ. Sleep to Remember. The Journal of Neuroscience. 2017; 37(3):457–463. https://doi.org/10.1523/JNEUROSCI.0297-16.2017, doi: 10.1523/JNEUROSCI.0297-16.2017.

Sara SJ, Bouret S. Orienting and reorienting: the locus coeruleus mediates cognition through arousal. Neuron. 2012; 76(1):130–141. https://doi.org/10.1016/j.neuron.2012.09.011, doi: 10.1016/j.neuron.2012.09.011.

Selinger JV, Kulagina NV, O’Shaughnessy TJ, Ma W, Pancrazio JJ. Methods for characterizing interspike intervals and identifying bursts in neuronal activity. J Neurosci Methods. 2007; 162(1-2):64–71. http://www.ncbi.nlm.nih.gov/pubmed/17258322, doi: 10.1016/j.jneumeth.2006.12.003.

Skelin I, Kilianski S, McNaughton BL. Hippocampal coupling with cortical and subcortical structures in the context of memory consolidation. Neurobiology of Learning and Memory. 2018; http://www.sciencedirect.com/science/article/pii/S107474271830090X, doi: 10.1016/j.nlm.2018.04.004.

Straube T, Frey JU. Involvement of beta-adrenergic receptors in protein synthesis-dependent late long-term potentiation (LTP) in the dentate gyrus of freely moving rats: the critical role of the LTP induction strength. Neuroscience. 2003; 119(2):473–9. http://www.ncbi.nlm.nih.gov/pubmed/12770561.

Swift KM, Gross BA, Frazer MA, Bauer DS, Clark KJD, Vazey EM, Aston-Jones G, Li Y, Pickering AE, Sara SJ, Poe GR. Abnormal locus coeruleus sleep activity alters sleep signatures of memory consolidation and impairs place cell stability and spatial memory. Current Biology. 2018; 28(22):3599–3609.e4. http://dx.doi.org/10.1016/j.cub.2018.09.054, doi: 10.1016/j.cub.2018.09.054.

Takahashi K, Kayama Y, Lin JS, Sakai K. Locus coeruleus neuronal activity during the sleep-waking cycle in mice. Neuroscience. 2010; 169(3):1115–26. http://www.ncbi.nlm.nih.gov/pubmed/20542093, doi: 10.1016/j.neuroscience.2010.06.009, takahashi, K Kayama, Y Lin, J S Sakai, K Neuroscience. 2010 Sep 1;169(3):1115–26. doi: 10.1016/j.neuroscience.2010.06.009. Epub 2010 Jun 11.

Tang W, Shin JD, Frank LM, Jadhav SP. Hippocampal-Prefrontal Reactivation during Learning Is Stronger in Awake Compared with Sleep States. The Journal of Neuroscience. 2017; 37(49):11789–11805. http://www.jneurosci.org/content/jneuro/37/49/11789.full.pdf, doi: 10.1523/jneurosci.2291-17.2017.

Totah NK, Neves RM, Panzeri S, Logothetis NK, Eschenko O. The Locus Coeruleus Is a Complex and Differentiated Neuromodulatory System. Neuron. 2018; 99(5):1055–1068 e6. https://www.ncbi.nlm.nih.gov/pubmed/30122373, doi: 10.1016/j.neuron.2018.07.037.

Ul Haq R, Liotta A, Kovacs R, Rösler A, Jarosch MJ, Heinemann U, Behrens CJ. Adrenergic modulation of sharp wave-ripple activity in rat hippocampal slices. Hippocampus. 2012; 22(3):516–533. https://doi.org/10.1002/hipo.20918, doi: 10.1002/hipo.20918.

Varela F, Lachaux JP, Rodriguez E, Martinerie J. The brainweb: Phase synchronization and large-scale integration. Nature Reviews Neuroscience. 2001; 2(4):229–239. https://doi.org/10.1038/35067550, doi: 10.1038/35067550.

Vazey EM, Aston-Jones G. Designer receptor manipulations reveal a role of the locus coeruleus noradrenergic system in isoflurane general anesthesia. Proceedings of the National Academy of Sciences of the United States of America. 2014; 111(10):3859–3864. https://doi.org/10.1073/pnas.1310025111, doi: 10.1073/pnas.1310025111, lC and ACC Tech paper.

Wang DV, Ikemoto S. Coordinated Interaction between Hippocampal Sharp-Wave Ripples and Anterior Cingulate Unit Activity. The Journal of Neuroscience. 2016; 36(41):10663–10672. http://www.jneurosci.org/content/jneuro/36/41/10663.full.pdf, doi: 10.1523/jneurosci.1042-16.2016.

Wang DV, Yau HJ, Broker CJ, Tsou JH, Bonci A, Ikemoto S. Mesopontine median raphe regulates hippocampal ripple oscillation and memory consolidation. Nat Neurosci. 2015; advance online publication. https://doi.org/10.1038/nn.3998, doi: 10.1038/nn.3998 http://www.nature.com/neuro/journal/vaop/ncurrent/abs/nn.3998.html##supplementary-information.

Yang M, Logothetis NK, Eschenko O. Occurrence of Hippocampal Ripples is Associated with Activity Suppression in the Mediodorsal Thalamic Nucleus. J Neurosci. 2019; 39(3):434–444. https://www.ncbi.nlm.nih.gov/pubmed/30459228 https://www.ncbi.nlm.nih.gov/pmc/articles/PMC6335745/pdf/zns434.pdf, doi: 10.1523/JNEUROSCI.2107-18.2018.

